# DNA methylation assigns sequencing reads to closely related bacterial strains when their sequence cannot

**DOI:** 10.64898/2026.11.09.765334

**Authors:** Kardokh Kakabra, Marcus J. Claesson

**Affiliations:** School of Microbiology, University College Cork, Cork, Ireland; APC Microbiome Ireland, University College Cork, Cork, Ireland

**Keywords:** bacterial methylome, restriction-modification, strain resolution, nanopore sequencing, long-read phasing, genomic epidemiology

## Abstract

Clinical and public-health questions often concern strains rather than species, yet co-resident strains of one species can be so closely related that a single long read carries no sequence difference between them. We measured how often this happens in 639 pairs of isolates of *Enterococcus faecium*, *Klebsiella pneumoniae*, *Listeria monocytogenes* and *Staphylococcus aureus* sequenced on one nanopore platform. Among the 297 pairs above 99% average nucleotide identity (ANI), 40.37% of read placements (a read counted against one strain’s genome) carried no sequence difference between the two strains, and above 99.9% ANI about three quarters did. Strains of one species also differ in the DNA motifs they methylate. Our method, Epilect, assigns single nanopore reads to strains from the basecaller’s methylation calls, given a methylation profile for each strain, and declines to call when the evidence is weak. In the pairs above 99% ANI it correctly assigned 21.8% of the read placements that sequence could not place, at a precision of 0.959. In computational mixtures of 65 pairs of distinct strains at or above 99.5% ANI that were predicted to differ in methylation, its per-read F1 score exceeded that of the best of three sequence-based read-phasing tools by 0.25 (95% CI 0.08 to 0.37); where equally related strains were predicted to share a methylome, the comparison reversed. DNA methylation therefore supplies strain evidence on reads whose sequence carries none, and whether it will do so for a given pair of strains can be estimated before sequencing.

**Impact Statement:** Genomic surveillance, transmission studies and microbiome research increasingly ask which of several co-resident strains carries a gene, a plasmid or a resistance determinant. Long reads are expected to answer such questions because one read can span a gene and its genomic context. This study shows that, for closely related strains of one species, a large share of long reads carries no sequence evidence of the strain they came from, which limits every method that reads only sequence. It shows that bacterial DNA methylation, which nanopore basecallers already report on every read, recovers part of that missing information with high precision, and that the pairs of strains for which this works can largely be identified beforehand from their lists of methylated motifs. The contribution is a measured limit for sequence-based read assignment and a second, independent source of strain evidence that applies within a defined range of strain pairs, rather than a general replacement for sequence-based phasing. The software, a test dataset and all analysed data are public, so laboratories that sequence isolates on nanopore platforms can check before sequencing a mixture whether methylation is likely to separate their strains.

**Data Summary:**

1. Epilect source code, documentation for installation and use, and a test dataset with the exact command and parameter settings (demo/): https://github.com/kardokhk/epilect (MIT licence). Version 0.2.1 is archived at Zenodo, https://doi.org/10.5281/zenodo.22856319; the concept DOI https://doi.org/10.5281/zenodo.22847418 resolves to the latest version. Analyses were run with the development code from which v0.2.0 was released. v0.2.0 corrected the motif-density count for palindromic motifs and added a read-length option for locus attribution and two memory options; Tables S33 and S55 were regenerated with it, and it changed no read assignment. v0.2.1 changed no code.
2. The 79-isolate panel: Dabernig-Heinz J *et al.* Nanopore based bacterial genotyping/surveillance in multi-center performance study. NCBI BioProject PRJNA1091452 (2024) [1, 2].
3. The co-resident isolate collection: Ürel H *et al.* Nanopore metagenomic sequencing links clinically relevant resistance determinants to pathogens. European Nucleotide Archive PRJEB108076 (2026) [3].
4. The *Streptococcus dysgalactiae* subsp. *equisimilis* methylome libraries: Schababerle T *et al.* Zenodo, https://doi.org/10.5281/zenodo.15555625, https://doi.org/10.5281/zenodo.15558488 and https://doi.org/10.5281/zenodo.15579791 (2025) [4].
5. REBASE allmeths.txt [5], retrieved 2026-09-09 (370,632 records).
6. Table S70 lists the BioSample and run accessions for every isolate and library analysed, under the identifiers used in this paper. Supplementary Tables S1 to S80 are provided as one Excel workbook, and Figs S1 to S9 in the Supplementary Material.

The authors confirm all supporting data, code and protocols have been provided within the article or through supplementary data files.

## Introduction

Many questions in clinical and public-health microbiology are asked of strains rather than species: which of two co-resident lineages carries a carbapenemase, whether an isolate from a patient is the one cultured last month, or whether a plasmid has moved between strains in one gut. Culture-enriched metagenomics found multi-strain gut colonisation by vancomycin-resistant *E. faecium* in 5 of 35 patients, diversity that isolate-based sequencing can miss [6]. Long-read sequencing makes such questions tractable in principle, because a single read can span a gene and its genomic context. Read-level phasing tools now separate co-resident lineages by the sequence differences between them [7–10], long reads can identify isolated or mixed strains of one species [11], and mixed samples can be deconvolved into lineages for genomic epidemiology [12–14]. All of these approaches need the distinguishing sequence to be present in the data, and at read level, on the read itself. One long-read method grouped strains separated by only a few mutations into a single group of siblings [11]. Conspecific isolates can differ by a handful of substitutions genome-wide, and homologous recombination exchanges sequence in stretches rather than one base at a time [15–17], so the differences that do exist can be concentrated in some regions and absent from long stretches of others. A read from such a stretch carries no sequence evidence of its strain, whatever its length.

Bacterial DNA carries a second layer of information. Most bacterial genomes examined carry methylated bases at short sequence motifs [18], and much of that methylation is the work of restriction-modification systems. In such a system a DNA methyltransferase methylates a short recognition motif throughout the cell’s own chromosome, while a partner restriction enzyme cuts the same motif where it is unmethylated, as on incoming phage DNA [19, 20]. Because the motif, rather than the individual site, determines what is methylated, most occurrences of a motif share one state, and the pattern is largely a property of the strain.

This pattern varies within species. Restriction-modification systems are frequently gained and lost and differ between lineages [19, 21, 22], so two isolates of one species can carry different methyltransferases and methylate different motifs. In a recent survey of 22 *Pseudolactococcus laudensis* genomes, nearly all strains had distinct methylation profiles, including strains isolated from the same dairy starter culture [23]. Not all methylation varies in this way: methyltransferases without a partner restriction enzyme are common [18], and many of these orphan enzymes are nearly as conserved within a genus as an average gene [22]. Where the set of methylated motifs, the methylome, does differ between two strains, it can be used to tell them apart.

Long-read platforms read methylation directly from the sequencing signal, with no bisulfite conversion and no separate library. Single-molecule real-time (SMRT) sequencing detects N6-methyladenine and N4-methylcytosine from the kinetics of DNA synthesis [18, 24, 25], nanopore sequencing detects these and 5-methylcytosine from the electrical signal of each molecule [26, 27], and current nanopore basecallers call all three on every read [28]. Nanopore methylome profiles were reproducible between replicate libraries for two *Streptococcus dysgalactiae* subsp. *equisimilis* strains at the three motifs tested [4]. The methylation state of each read is therefore available at no extra experimental cost whenever DNA is sequenced without amplification. Existing tools use these calls at the level of genomes and assemblies, pooling them across the reads covering each site to discover methylated motifs, describe a methylome, group metagenomic contigs or link plasmids to their hosts [27, 29, 30]. Single reads can also be grouped: SMRT reads from co-sequenced strains of one species were clustered into strain-level bins by their methylation profiles [31, 32], and nanopore reads and contigs carrying resistance genes were recently linked to their host species in mock communities and rectal swabs by methylation similarity [3].

Supervised assignment at strain level has, to our knowledge, not been reported: deciding which of two strains of one species a single nanopore read came from, using a methylation profile measured beforehand for each strain, with a measured error rate, and knowing before sequencing whether this is possible for a given pair. Its value depends on a quantity that has not been measured directly, namely how often reads from two closely related strains carry no sequence difference at all. Here we measure that fraction on a panel of isolates of four species, show how many of those reads methylation can assign, identify the pairs of strains in which methylation outperformed sequence-based tools, and test how well those pairs can be identified from motif lists before sequencing. The method, Epilect, compares the methylation calls on each read with the pattern expected for each strain, combines reads into a call for a locus, and reports when the evidence is too weak to decide.

## Methods

### Datasets

The panel is 79 bacterial isolates sequenced on Oxford Nanopore R10.4.1 by Dabernig-Heinz *et al.* [1, 2] (BioProject PRJNA1091452), 77 of them from four species, *E. faecium*, *K. pneumoniae*, *L. monocytogenes* and *S. aureus*. Two isolates form no within-species pair: one is *Enterococcus faecalis* and one has an unsupported label, its closest reference being *Staphylococcus simulans* at 89.67% ANI. Pairwise ANI was computed with skani [33], and species assignment used the 95 to 96% ANI boundary [34]. Isolates were grouped into clones by single linkage at 99.99% ANI, giving 50 clones from the 77 isolates that enter pairs. Within-species pairs that are not near-clonal number 639. Of these, 620 are within-species by ANI against type strains: the 19 pairs formed with isolate Kp17 are between-species, because Kp17 is *Klebsiella quasipneumoniae* subsp. *similipneumoniae* (99.14% ANI to the type strain at 91.9% aligned fraction, against 94.01% to *K. pneumoniae*). Per-species and per-band statements use the 620 within-species pairs; including the Kp17 pairs, the lowest band is 0.41% over 110 pairs (Table S71). The 91 *L. monocytogenes* pairs below 95% ANI fall between two within-species clusters whose membership matches curated lineage I and lineage II sequence types exactly [35, 36], and they remain conspecific under NCBI Taxonomy [37] and LPSN [38], which treat the species as one. The classification is convention-dependent: GTDB release R232 [39] assigns genomes of this species to more than one species-level cluster, placing the type strain and the lineage I reference F2365 in different clusters. We did not establish that the GTDB partition corresponds to the lineage I and lineage II division, and we do not claim it.

The controlled two-strain mixture used two *Streptococcus dysgalactiae* subsp. *equisimilis* isolates, UT9728 and UT10237, from the methylome libraries of Schababerle *et al.* [4]. The co-resident isolate collection, which provided the two-strain library RS5_barcode12, specimen RS8 and the replicate colony picks, is European Nucleotide Archive study PRJEB108076 from Ürel *et al.* [3]. Table S70 links every isolate and library identifier used here to its BioSample and run accessions.

### Overview of Epilect

Epilect assigns a single long read to one of two or more co-resident strains using the methylation calls that the basecaller has already attached to the read. It needs no assembly or binning of the sample. It does need, for each strain, a description of that strain’s methylome: the motifs the strain methylates and how often a call at each motif is methylated. This description is the method’s principal precondition, and it must come from a reference methylome, from an isolate of the strain, or from motif discovery.

#### Scoring a read

For each motif, a read contributes the number of called sites and the number called methylated. Epilect computes how likely those counts are under each strain’s methylome, and the difference between the two log-likelihoods is the read’s evidence: its sign says which strain explains the read better, and its size says by how much (Fig. 1a to c). Calls are counted at their motif, so a methylated cytosine in CCWGG and one elsewhere are never pooled. Before scoring, a call is kept only if the four bases flanking it in the read agree with the reference, which removes calls made on misread sequence; we refer to this as the fidelity filter. A read is assigned when it carries at least one informative site and its evidence exceeds 1.1 natural-logarithm units (nats) in either direction; otherwise Epilect declines and records why. These default settings were selected under familywise control of multiple testing (Fig. S9; Tables S56 to S58).

**Fig. 1.**
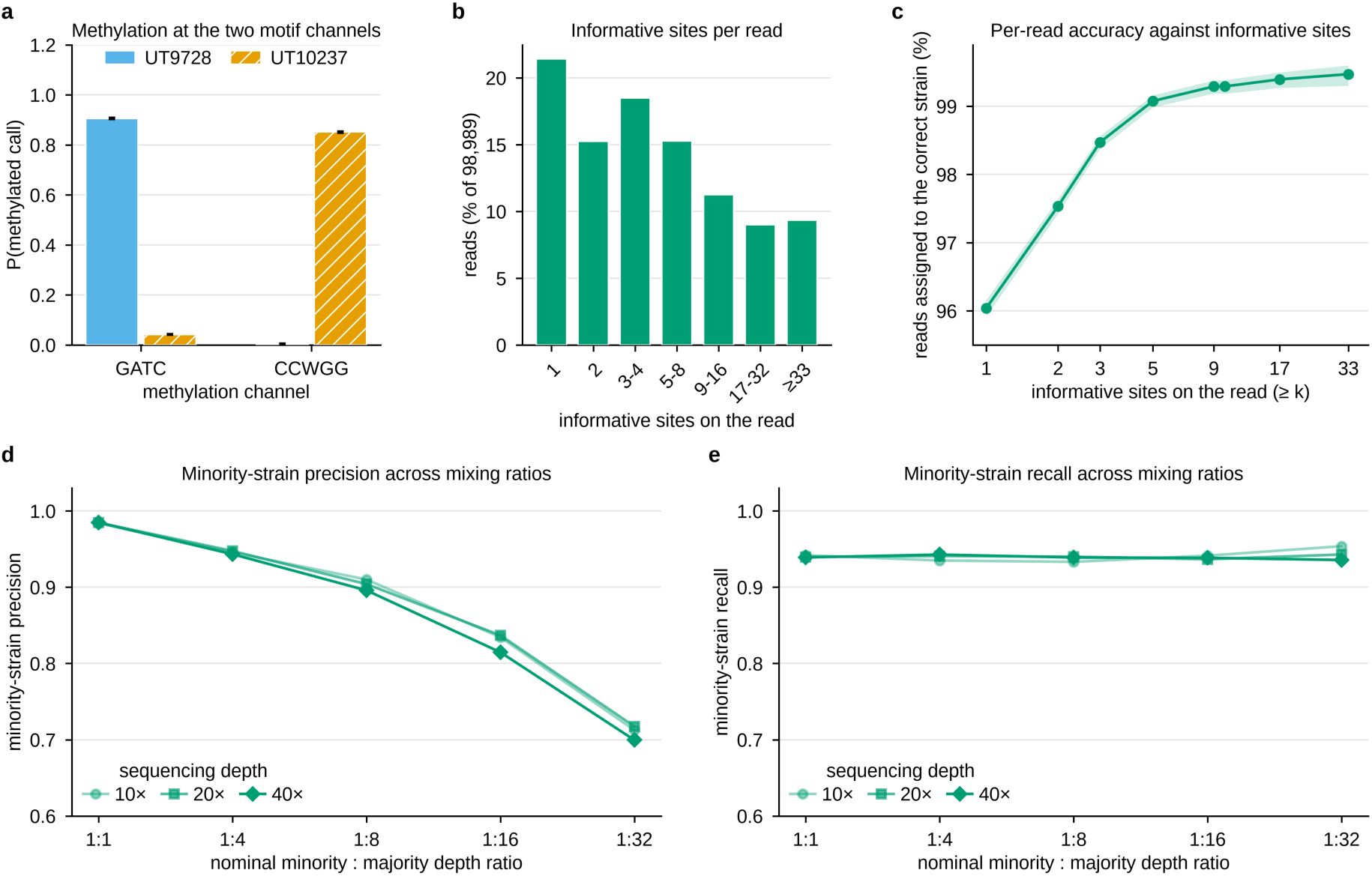
Assigning single reads to a strain from methylation alone. (a) Methylated fraction at the two motifs that separate the two *S. dysgalactiae* subsp. *equisimilis* isolates, measured on each isolate alone; error bars are 95% percentile intervals from a bootstrap resampling reads, not calls (2,000 replicates, seed 20260913). (b) Informative motif sites per read over 98,989 reads from the pure isolates. (c) Accuracy among reads carrying at least *k* informative sites, with 95% Wilson intervals; accuracy is the best-scoring strain with no threshold, no site minimum and no fidelity filter. (d, e) Minority-strain precision and recall in computational mixtures at three sequencing depths and five ratios, at default settings (fidelity filter on, evidence above 1.1 nats, at least one informative site). Every mixture result assumes a supplied methylome for each strain. See also Fig. S1.

#### From reads to a locus

Epilect combines the reads covering a locus into a single score and calls the locus only when that score passes a threshold. The score is capped both per read and in the number of reads it counts. Every read at a locus comes from one library, so without the cap a small library-wide bias would grow with depth until any locus appeared confidently called.

#### Predicting whether two strains can be separated

Whether one read can tell two strains apart depends on their methylomes and on read length, and Epilect computes it before any read is classified. From the two strains’ lists of methylated motifs it takes the motifs that only one strain methylates and estimates how many of their sites a read of a given length is expected to carry, with three sites per read as the threshold. The result places each pair in one of four resolvability classes: identical methylomes, where methylation carries no distinguishing signal; unshared motifs dense enough in both strains that reads of either can be assigned singly (both directions); dense enough in one strain only (one direction); or only sparse unshared motifs, so that single reads carry few informative sites and a confident call generally needs several reads (coverage only). The prediction needs motif lists and a read length but no measured methylation, so it can be made before sequencing.

### Sequence channel and the two definitions of sequence-blindness

Each ordered pair of genomes was aligned with one identical command, minimap2 [40] -cx asm20 --cs piped to paftools.js call -l 1000 -L 1000, for all 1,408 asm20 ordered-pair alignments, with nothing tuned per pair, per species or per class. Reads were mapped with minimap2 -y -ax map-ont. A read placement is one read against one strain’s reference. Each pair was analysed against both strains’ genomes, so a read contributes one placement per orientation, and an isolate’s reads recur in every pair it belongs to; placements are therefore not distinct reads, and every fraction reported for the panel is a fraction of read placements. A placement is sequence-blind under the permissive definition when no discriminating difference falls in its reference span, and under the core-restricted definition, used throughout the Results, when in addition at least 99% of that span lies inside the one-to-one alignment core. The two differ by 439,580 placements, of which 416,001 have no one-to-one alignment at all; there, presence and absence of sequence is itself evidence, and a sequence method has maximal rather than zero power. Under the permissive definition the sequence-blind fraction above 99% ANI is 48.50% rather than 40.37%, and the five ANI strata give 4.28%, 8.53%, 48.50%, 72.83% and 81.28% (Table S71). Over all 639 pairs the permissive fraction is 30.16% with minimap2 asm5 against 26.20% with asm20. On the permissive definition, Epilect assigned 23.57% of sequence-blind placements over all 639 pairs at a precision of 0.953 (Tables S6 and S72).

### Methylation channel and the classifier

Modified-base calls were produced with dorado [28] and extracted per call with modkit [41]. Each strain is described by an emission model: for every methylated motif, the probability that a call at an occurrence of that motif is methylated in that strain. Under a Bernoulli model with calls independent within a motif channel and channels independent across the read, the log-likelihood of a read under strain *s* is the sum over channels of *m* log *p*s + (*n* − *m*) log(1 − *p*s), where *n* is the number of admitted calls and *m* the number called methylated, and the read’s evidence is the difference between the two strains’ log-likelihoods. With more than two strains the statistic is the margin between the best strain and the second best. Emission probabilities are clamped away from zero and one, which bounds the influence of any single call. Calls are routed to channels by the motif annotation of the caller’s own output rather than by canonical base; a site matching more than one declared motif is assigned to the first declared and counted separately. A call is admitted when the caller did not flag it, when its motif is a declared channel, and when all four flanking bases of the caller’s five-base context agree between the reference and the read (the fidelity filter). The centre base is excluded, because it is the base whose modification state is in question. A read the filter empties is retained with an empty record, so it remains in the mapped total and contributes to no numerator. The default is the fidelity filter on, |evidence| > 1.1 nats and at least one informative site, selected under a familywise max-|z| multiplicity policy over 110 stratum-level tests per candidate setting (170 with read-length strata added, which selected the same setting) (Fig. S9; Tables S56 to S61); a setting with the filter off, |evidence| > 1.0 and at least two informative sites is available by flag and is the setting at which Fig. 2c was measured. Three further options are implemented and disabled by default because none improved the result it was designed to improve: per-call soft evidence in place of a hard call, a beta-binomial in place of the binomial, and masking of phase-variable loci.

**Fig. 2.**
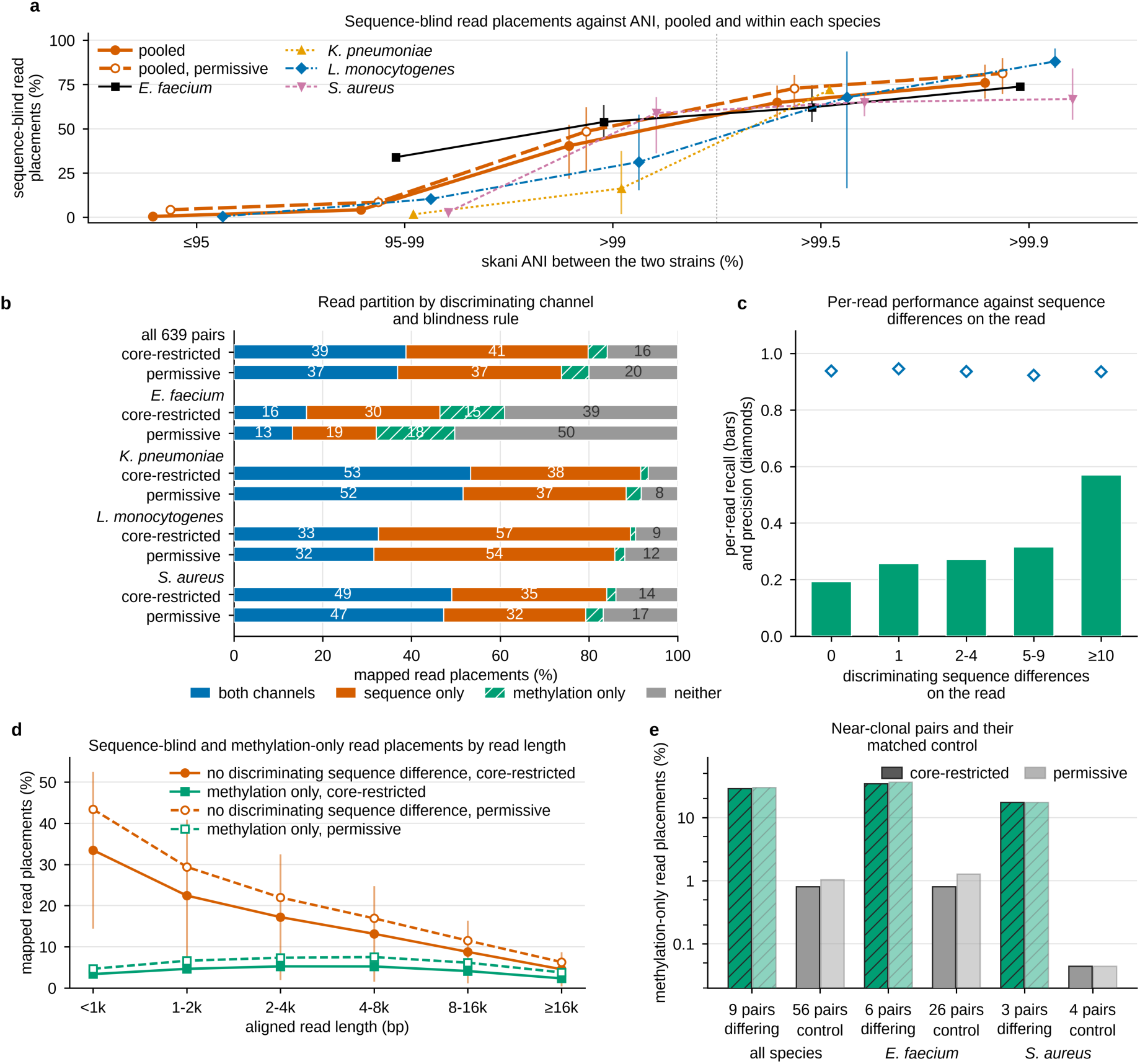
Read placements that carry no sequence difference between two strains. A read placement is one read counted against one strain’s genome. Panel (a) covers the 620 within-species pairs; panels (b) to (d) cover the 639 pairs of the 79-isolate panel and the 7,341,853 read placements mapped across them; panel (e) covers a separate set of 65 pairs of isolates from one clone. Both definitions of sequence-blindness, core-restricted and permissive, are shown (Methods). (a) Sequence-blind read placements against the pair’s ANI, pooled and within each species; three separate bands and two nested thresholds, divided by the dotted rule. Bars are dyadic cluster bootstrap intervals over clones, absent where fewer than six clones remain. (b) Read placements partitioned by which source of evidence can distinguish the two strains, pooled and by species. (c) Per-read recall and precision against the number of sequence differences a read carries, measured without the fidelity filter and with a two-site minimum. (d) Sequence-blind and methylation-only read placements by aligned read length. (e) Read placements assigned from methylation alone in the nine pairs from one clone that carry a methylome difference, beside 56 identically filtered pairs that carry none; no interval is given for the nine, which come from two clones. See also Fig. S2.

### Locus statistic and resolvability classes

The locus score is Λ = min(*n*, 10) × the mean per-read evidence clipped at ±5 nats, and a locus is called when |Λ| reaches the threshold (10 in all results). Clipping prevents one atypical read from dominating, and capping the effective read count bounds the score and makes its ceiling available in closed form. The negative-control contrast between capped and uncapped statistics holds at both classifier settings; values in the Results are at the default. Resolvability classes use the expected density of unshared-motif sites per read computed from IUPAC degeneracy, with each palindromic site counted once, against a threshold of three sites per read. The panel classes and the released tool identify palindromes by comparing a motif with its reverse complement; the REBASE analysis uses REBASE’s strand annotation, and the two rules disagree for 405 of 3,443 motifs. The panel classes and the REBASE analysis sum density over each strain’s unshared motifs. The pair_tier function of the released tool, and the RS8 read-length prediction made with it, use instead the densest single unshared motif, a more conservative rule that agreed less well with read assignment (Results; Table S80).

#### Testing the predicted class against read assignment

Each pair’s observed class was built only from Epilect’s assignments of the held-out reads of its two isolates at default settings, with each strain’s genome serving as reference in turn; no motif list, motif density or methylation level entered it. A strain counted as separable from single reads when at least half of its mapped reads were assigned to it and at least 90% of the reads assigned to it were its own; precision was not evaluated below 20 assigned reads. A pair carried no signal when, for both strains, the share of mapped reads assigned correctly minus the share assigned incorrectly was at most 0.10, the 99th percentile of that quantity over the 56 near-clonal pairs with identical methylomes, computed before any other pair’s outcome was read. A pair with both strains separable was observed separable in both directions, one with one strain separable in one direction, and any other pair at coverage level. No pair was excluded; every direction had at least 1,282 mapped reads. The thresholds and the primary statistic, a linearly weighted κ over the four ordered classes with a dyadic cluster bootstrap over clones, were fixed in a written plan before any agreement was computed, except that the recall threshold of 0.50 was set after the outcomes of the nine near-clonal pairs with differing methylomes had been seen; thresholds of 0.40 and 0.60 and the alternative classifier setting are therefore reported as stability analyses (Table S80). Three couplings remain between prediction and outcome. The default settings were selected on these pairs’ outcomes; the motif lists behind both the prediction and the classifier were discovered from all reads of each isolate; and each isolate was classified with its own methylation model. The observed resolving power is therefore an upper limit on what the classifier would achieve for strains not used in this way.

### Recombination analysis

Per species, a core pseudo-alignment of the panel isolates was built and recombination inferred with Gubbins [42]; ClonalFrameML [43] was run on the resulting tree. Annotation for the mobile-element mask used Prokka [44] and PhiSpy [45]. Three criteria were fixed before any result was inspected: a majority of between-clone substitutions inside inferred blocks; a density-independent allele-sharing test against unrelated lineages; and persistence after masking prophage, mobile-element coding sequences, repeats and contig ends. The criteria were pre-specified; the numeric thresholds, the bootstrap lower-bound rule and the contig-edge mask were not. ClonalFrameML is not an independent detector here, because it consumed the Gubbins tree.

### Benchmark against sequence-based tools

Comparators were Floria [7] at its default (–snp-count-filter 100) and at --snp-count-filter 2, devider [9] at its default and at --min-cov 1 --min-abund 0.1, and HairSplitter [10] at its default; in each mixture the comparator is the highest-scoring of the five. Strainy [8] was run as a sixth comparator on a pre-declared subsample and is not reproducible run to run, because metaFlye pins no seed in this invocation. Each benchmark pairing is two panel isolates with one serving as the reference. Mixtures were built from held-out reads of each isolate (80,000 per isolate, 40-fold total, subsample seed 11) at 1:1, 1:4 and 1:16 with the non-reference isolate as the minority, and 1:16 with the reference isolate as the minority. One offered read set was used per mixture, the primary non-supplementary mapped records, and every tool received that alignment or the reads made from it. Each read contributes one row and takes the label of its longest part; F1 is computed over every offered read, an undefined F1 counts as zero, and clusters are mapped to strains by reference anchoring, which gives no tool read-level truth (Fig. S7; Tables S44 to S46). Discriminating single-nucleotide differences come from assembly-to-assembly alignment (minimap2 asm5 with paftools.js call), never from the test reads, and each strain’s methylome from reads excluded from every mixture. The target regime was defined after a first benchmark of 52 pairings (208 mixtures; Tables S38 to S43), 12 of which fall in it, from two properties known before sequencing: an ANI band (at or above 99.99%; 99.50 to 99.99%; 98.50 to 99.50%; below 98.50%) and the predicted resolvability class. The remaining pairings were selected by those two properties alone, before any tool was run on them: every panel pairing in the target regime, and an off-target set matched on species and, within species, on clone, ties broken by a deterministic sort on pairing identifier (Table S73). A sensitivity analysis runs every tool, Epilect included, at its own published defaults. Standard phasing metrics on the controlled mixture, a masked-variant sweep, an *S. aureus* pair, a three-strain mixture, block orientation accuracy and per-read label accuracy are reported in Fig. S8 and Tables S62 to S67, with whatshap [46] for phasing metrics and Flye [47] for assembly. Measured wall time per mixture, with 16 invocations sharing one node, was 1.5 to 4.5 times that of Floria and devider for Epilect, and peak memory 4.1 to 5.0 times (controlled mixture) and 6.3 to 7.9 times (panel); Epilect ran in 0.41 and 0.39 times the wall time of HairSplitter and in 0.06 times that of Strainy’s phasing step (Table S74). These tools answer a different question from Epilect, and the ratios are reported as measurements rather than as a ranking.

### Run time and memory

Cost was measured on one MeluXina 128-core compute node with one invocation at a time, three replicates per point, at eight threads unless stated, for seven samples of *S. dysgalactiae* subsp. *equisimilis*, *S. aureus* and *Escherichia coli* (0.0116 to 0.5953 Gb of aligned read sequence). Assigning the reads of one sample took 2.05 s plus 101.8 s per gigabase of aligned read sequence, so a 5 Mb genome at 40-fold coverage takes about 22 s and 3 to 9 GB of memory (Table S74). Wall time is the median of three replicates; no interval is formed, and the linear fit (R² = 0.997) describes these seven points rather than predicting with stated coverage. Nearly all of the time and memory is spent in modkit, which extracts the per-call data from the aligned reads; reading modified-base tags directly with pysam was measured and rejected, because decoding them alone took longer than the whole modkit extraction. Peak memory per gigabase was 36.7 to 43.6 GB on a single closed chromosome, 27.8 on three contigs and 14.8 on 168, so a memory estimate needs the contig count as well as the input size. Memory can be exchanged for time with identical output: -- modkit-interval-size 10000 reduces peak memory 6.2-to 6.6-fold for 1.34 to 1.56 times the wall time (30000 gives 2.6-to 2.9-fold for 1.09 to 1.16 times), and --modkit-ignore-index reduces it 16.5-to 36.7-fold, to 0.10 to 0.24 GB, for 3.19 to 4.46 times, on both classify and attribute. For classify, output was bit-identical across all settings in 33 read-by-read comparisons over 1,588,180 reads; for attribute, the settings were checked only on the bundled demonstration data. At 32 threads assignment ran 1.7 to 2.0 times faster for 2.2 to 3.2 times the peak memory. Attributing 251 loci on an 8.04 Mb sample took 93.02 s and 9.43 GB, and the check that decides which loci can be attributed took 1.54 s.

### Software versions

epilect development code from which v0.2.0 was released (Data Summary) on Python 3.12.14; dorado 2.1.2 [28], ont-modkit 0.6.4 [41], samtools and bcftools 1.23.1 [48], whatshap 2.8 [46], Flye 2.9.6 [47], nanomotif 1.1.2 [29], minimap2 2.30 [40], skani [33], Gubbins 3.4.3 [42], ClonalFrameML 1.20 [43], Prokka 1.15.6 [44], PhiSpy 5.0.10 [45], AMRFinderPlus 4.2.7 [49], sequence typing against PubMLST [50] with blastn 2.15.0+ [51], and pysam 0.24.1. REBASE allmeths.txt carries no internal version; its FTP modification date is 2024-03-29 and it was retrieved on 2026-09-09, giving 370,632 records. Tables S68 and S69 list the software, versions and citations. Nanomotif was rerun with version 1.3.0 on the same inputs and parameters, and every value quoted here was unchanged. Seeds are stated with each resampling procedure.

### Statistical analysis

#### The unit of analysis is the clone

The 79 isolates collapse to 50 clones at 99.99% ANI, so pairs and reads are not independent observations, and every interval on the panel resamples clones rather than isolates, pairs or reads. Two clone-level intervals are computed. The dyadic cluster bootstrap resamples clones with replacement and takes all pairs among the resampled multiset (2,000 replicates, seed 20260913). The paired leave-one-clone-out jackknife deletes every pair or mixture involving the deleted clone and is quoted as the estimate plus or minus 1.96 standard errors. Where both were computed, the text quotes the bootstrap, because clone deletion on this panel is unbalanced and the jackknife assumes balanced deletion: one *E. faecium* clone touches 43% of the target-regime mixtures of distinct strains, and above 99% ANI deleting clone Ef28 removes 84 of 297 pairs (sequence-blind fraction 36.06%) while deleting Ef40 moves the fraction furthest, to 35.19% on 70 pairs. Both intervals are given in Tables S71 and S72; for the sequence-blind fraction above 99% ANI the jackknife gives 22.75% to 58.00% against the bootstrap’s 21.85% to 52.26%. The jackknife is not constrained to the range of a proportion, so some lower bounds fall below zero; these are reported as computed. An interval is not formed where fewer than six clones remain.

#### Where no interval is given, the reason is stated

No interval is quoted for the nine near-clonal pairs carrying a methylome difference, which sit in two clone clusters; for the 94.6% recombination-block fraction, whose signal rests on about three inferred ancestral events; for the ratio of median F1 in the benchmark, whose denominator is close to zero; for any single-species estimate in the benchmark target regime, each of which rests on fewer than six clones; or for single-specimen results. The share of placements that never reach the classifier is quoted as a count and share, because it varies by more than an order of magnitude between clones.

#### Equivalence of methylome and sequence divergence

An equivalence test at |ρ| < 0.30 is uninformative on this panel in either direction: with 8 to 14 clones per species, the power to declare equivalence when the truth is zero is 0.08 to 0.67 under a structure-preserving null and 0.00 to 0.35 under a clone random-effect null.

#### Decomposition of the ANI ladder

Holding the four species at their above-99% levels and reweighting them to the species mix of the pairs above 99.9% predicts 48.54% against the 75.93% measured, so the change of mix accounts for 0.0817 of that step’s 0.3555, a share of 23.0% (Table S71). The decomposition is partial and non-orthogonal, because the above-99% baseline contains the above-99.9% pairs, and it carries no interaction term.

## Results

### A large share of reads from closely related strains carries no sequence difference

To measure how often sequence alone cannot place a read, we formed every pair of panel isolates labelled as one species that were not copies of a single clone: 639 pairs, of which 620 are within one species by genome comparison (Methods). For each pair we aligned the two genomes and asked, of every mapped read, whether any sequence difference between the two strains fell within the stretch of genome it covered. A read placement was sequence-blind when no difference fell in its span and at least 99% of the span lay in the part of the two genomes that aligns one-to-one, the stricter of the two definitions we examined (Methods; Tables S4 to S7). The sequence-blind fraction rose with relatedness, from 0.49% of read placements in pairs at 95% ANI or below to 40.37% above 99% (95% CI 21.85 to 52.26) and 75.93% above 99.9% (Fig. 2a; Table S71). Every step of this ladder separated from zero. ANI was not manipulated, so the fraction is associated with relatedness rather than shown to be caused by it, and each value describes this panel’s pairs rather than a randomly drawn pair of strains.

The rise is visible inside each species, so it is not merely a change in which species occupy which band, although the within-species evidence is weaker than the pooled: of the twelve steps the four species can form, eight separated from zero, and the four that did not all lie at the top of the ladder, where each species contributes fewest pairs (Fig. 2a). No interval around the pooled value covers the spread between the species, so the per-species values are the ones to read. The species mix also accounts for part of the pooled rise, 23.0% of the step above 99.9% ANI, in a partial decomposition (Methods; Table S71).

In *E. faecium*, 94.6% of the substitutions separating a pair fell inside recombination blocks inferred on the branches separating them. Substitutions in blocks also carried an allele found in an unrelated lineage more often than background substitutions did, and both results held after prophage, mobile elements, repeats and contig ends were masked. The signal rests on about three inferred ancestral events, so we give no interval for the 94.6% and claim recombination for this species only. In *K. pneumoniae*, *L. monocytogenes* and *S. aureus* the substitutions are also clustered, but neither test met the criteria fixed in advance, and we name no mechanism for them.

### Methylation assigns reads that sequence cannot

If 40% of read placements between closely related strains carry no sequence difference, the question is whether methylation can place them. We first established how single reads behave in a controlled mixture and then measured performance on the sequence-blind reads of the panel.

The controlled mixture was built from two *S. dysgalactiae* subsp. *equisimilis* isolates [4] whose methylomes differ at two motifs. Measured on each isolate alone, UT9728 methylated GATC at 0.9057 and CCWGG at 0.0047, whereas UT10237 methylated GATC at 0.0418 and CCWGG at 0.8516 (Fig. 1a; Table S1). The intervals shown there resample reads rather than calls, because calls on one read are not independent.

Most reads carry little evidence: of 98,989 reads from the pure isolates with at least one usable call, 21.41% carried a single informative site (Fig. 1b). Nevertheless, the better-scoring strain, taken with no threshold, site minimum or fidelity filter, was the correct one for 96.04% of reads (95% CI 95.92 to 96.16), rising to 99.29% among reads carrying nine or more informative sites (Fig. 1c; Table S10).

We then mixed reads of the two isolates computationally at five ratios, from 1:1 to one part in 32, and three sequencing depths, 10-, 20- and 40-fold. Recall of the minority strain, the share of its reads assigned to it, stayed high at every ratio and depth, whereas precision, the share of reads assigned to it that truly came from it, fell as the minority grew rarer (Fig. 1d, e; Fig. S1; Tables S16 and S17). The fidelity filter holds precision up in skewed mixtures: at 40-fold and 1:32, switching the filter on alone, at |evidence| > 1.0 nats and a two-site minimum, raised minority-strain precision from 0.51 to 0.71 (Table S75). Across the panel’s pairs, its effect on the minority strain’s F1 score, which combines precision and recall on a scale from 0 to 1, ran from a small loss at 1:1 to a gain of 0.0505 (95% CI 0.0368 to 0.0641) at 1:32 (Fig. S9b; Table S58).

All of these results assume that each strain’s methylome is known, and that assumption is strict. When the methylomes had instead to be discovered from a two-strain mixture mapped to a single reference, without assembly or binning, discovery failed at every ratio tested: a usable model was produced in only 5 of 270 mixtures, and relative to supplying the true motifs, minority-strain F1 fell by 0.306 (95% CI 0.091 to 0.520) at 1:1 (Table S76). Where an assembly separates the two strains into contigs, the same command does recover a minority strain’s motifs, so the limit lies in the single-reference setting rather than in the discovery algorithm (Tables S19 and S20).

On the panel the test is sharper, because a sequence-blind read offers a sequence-based method nothing to use. Among the 297 pairs above 99% ANI, Epilect assigned 21.8% of sequence-blind read placements correctly (298,073 of 1,365,407) at a precision of 0.959, and over all 639 pairs it assigned 21.42% at the same precision (Tables S8 and S72). No sequence-based tool was run for this comparison, because on these reads the discriminating sequence is absent by definition.

A contrast within the panel indicates that these assignments come from methylation. Nine pairs of isolates from a single clone happen to differ in methylome, whereas 56 otherwise identically filtered pairs from a single clone do not. In the nine, Epilect assigned 28.91% of read placements from methylation alone at a precision of 0.960, against 0.81% in the 56. The nine pairs come from only two clones, so an interval over clones would reflect little more than whether those two are drawn. The contrast instead holds separately within both species (Fig. 2e; Tables S23 and S24). Methylation’s precision on sequence-blind reads did not fall as strains became more similar. Within each species, precision showed no trend with ANI, every slope having an interval that covered zero, and it stood between 0.920 and 0.965 across the four species (Table S72). Recall likewise showed no trend in three species, and in *S. aureus* it fell as strains became more similar.

The two sources of evidence are not complementary, and this bounds what methylation can do. Recall rose with the number of sequence differences a read carried, from 0.170 on sequence-blind read placements (0.193 under the permissive definition shown in Fig. 2c) to 0.570 on those carrying ten or more (Fig. 2b, c; Tables S8 and S21; measured without the fidelity filter and with a two-site minimum). Methylation is therefore weakest on precisely the reads for which it is the only evidence. Read length narrows its role further, since the sequence-blind fraction fell from 33.48% of read placements below 1 kb to 4.55% above 16 kb (Fig. 2d; Tables S25 and S26). Finally, 14.47% of read placements never reached the classifier, because they carried no methylation call at all or were emptied by the fidelity filter (Table S72).

### Motif lists predict whether single reads can separate two strains

Not every pair of strains differs in methylation, so a laboratory needs to know in advance whether methylation will separate its strains. The resolvability class answers that from motif lists (Methods), and we tested it against what the panel’s reads delivered.

Predicted and observed classes agreed for 333 of the 639 pairs (52.1%; linearly weighted κ = 0.57, 95% CI 0.39 to 0.71 over 50 clones; Fig. 3c; Table S27). Most of the disagreement lay between the two single-read classes: 200 of the 242 pairs predicted separable in one direction were separable in both, consistent with a motif methylated in only one strain marking the reads of both, methylated on one and unmethylated on the other. With the two single-read classes merged, the prediction placed 533 of the 639 pairs correctly among no signal, coverage-level signal and separation from single reads (83.4%; κ = 0.79, 0.61 to 0.91). None of the 132 pairs predicted to have identical methylomes was separable from single reads, although 39 of them (29.5%) carried a coverage-level signal. The three-level agreement held at every threshold examined but not in every species: it was lowest in *K. pneumoniae* (κ = 0.58, 0.13 to 0.92), where 57% of pairs were separated better than predicted (Fig. S4; Table S80).

**Fig. 3.**
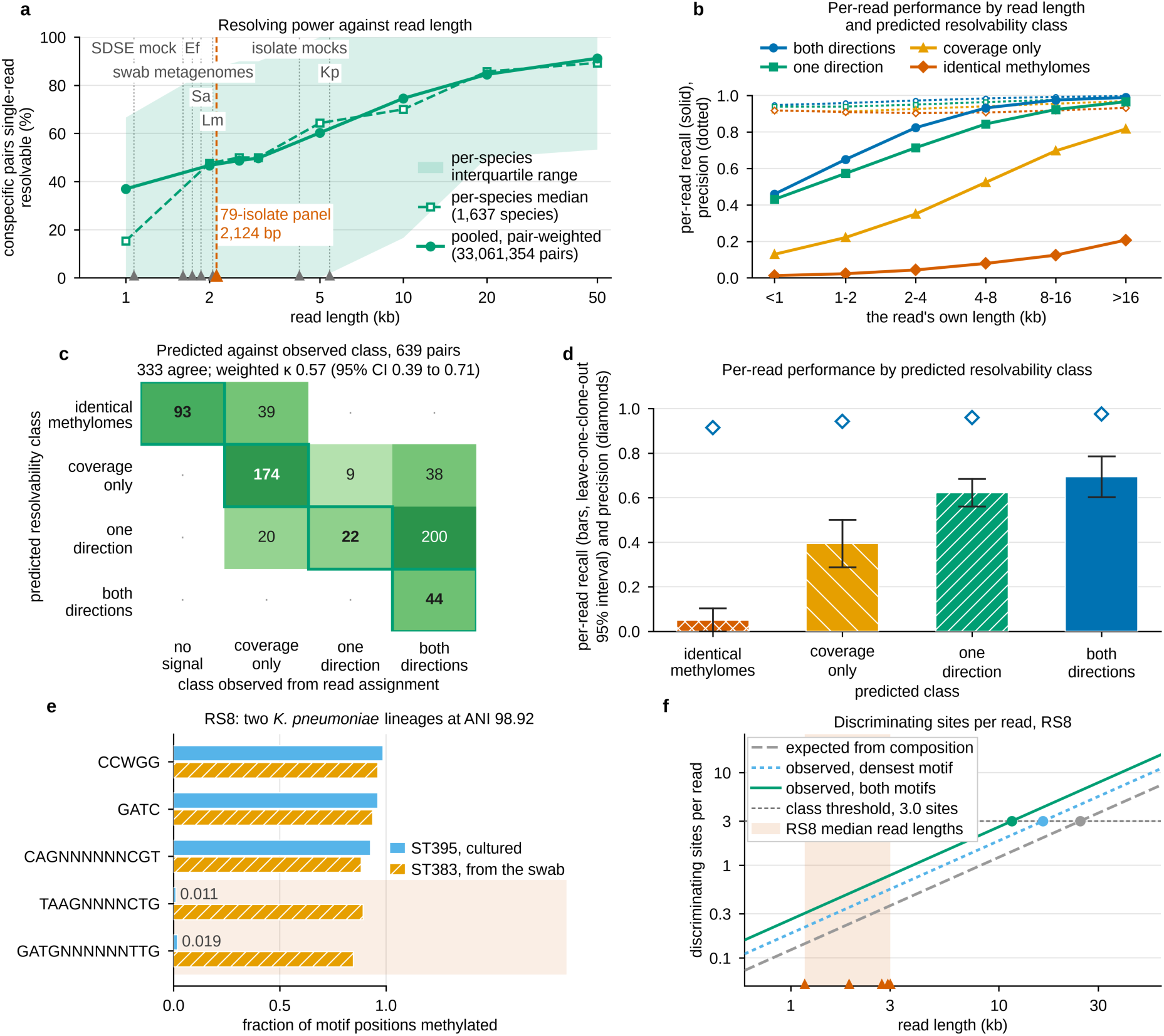
Predicting before sequencing whether single reads can separate two strains. (a) Fraction of pairs of one species in REBASE predicted separable from single reads, against read length, pooled over 33,061,354 pairs and as the median over 1,637 species with the interquartile range shaded. Dotted rules mark each dataset’s read length, measured identically for all: the median over primary aligned reads per run, then the median of per-run medians. (b) Per-read recall and precision against the read’s own length, by predicted resolvability class. (c) Predicted class against the class observed from read assignment at default settings, for 639 pairs; shading darkens with the logarithm of the pair count, a dot marks a combination with no pair, and the outlined cells with bold counts form the diagonal, where the classes agree: 333 pairs lie on it, 286 above it (separated better than predicted) and 20 below it (linearly weighted κ 0.57, 95% CI 0.39 to 0.71). The observed class “no signal” corresponds to the predicted class of identical methylomes. (d) Per-read recall by predicted class, with leave-one-clone-out intervals over 50 clones. (e, f) Specimen RS8, whose two *K. pneumoniae* lineages are at 98.92% ANI. See also Figs S3 and S4.

Where the prediction erred, it more often under-stated resolving power than over-stated it: 286 pairs (44.8%) were separated better than predicted and 20 (3.1%) worse, all 20 predicted separable in one direction and observed at coverage level. The size of this asymmetry depended on the recall threshold that defines separability, from 57.0% against 0.3% at a threshold of 0.40 to 24.1% against 14.2% at 0.60, and the lower bound of the four-class κ reached the 0.40 set in advance as support at 0.60 and at the alternative classifier setting, but not at 0.40 or 0.50 (Table S80). Because the observed resolving power is an upper limit (Methods), under-prediction is likely to be less frequent, and over-prediction more frequent, for strains whose reads were not used to tune the classifier. The rule as implemented in the released tool, which takes the densest single unshared motif rather than summing over unshared motifs, agreed less well (κ = 0.38, 0.19 to 0.56) and under-predicted in 53.7% of pairs (Table S80).

The class also predicted how well single reads would be assigned, although it separated them into three levels rather than four. Per-read recall rose from 0.050 where methylomes are identical to 0.394 where only coverage separates the strains, 0.622 in one direction and 0.694 in both (Fig. 3d). The first two steps separated from zero, whereas the step from one direction to both did not. Within every class, recall rose steeply with the read’s own length, in the one-direction class from 0.431 below 1 kb to 0.966 above 16 kb, while precision stayed above 0.90 in all 24 combinations of class and length (Fig. 3b).

ANI is no substitute for this prediction, because the methylome difference between two strains is not a simple function of their sequence difference. Among pairs above 99% ANI the two were associated in *S. aureus* (Spearman ρ = −0.82, 95% CI −0.98 to −0.38; 34 pairs, 12 clones), whereas the pooled estimate straddled zero (−0.29, −0.65 to 0.06; 297 pairs, 47 clones; Fig. S2; Tables S9 and S77).

One patient specimen gave a test outside the panel. Specimen RS8 carried two *K. pneumoniae* lineages, a cultured ST395 isolate and an ST383 lineage assembled from the same rectal swab, at 98.92% ANI. Reads would need to reach 24,576 bp to carry three distinguishing sites, against median read lengths of 1,168 to 3,011 bp in the specimen’s five libraries, so single reads were predicted not to separate the lineages (Fig. 3e, f; Table S33). This is one specimen and settles nothing alone. Because the prediction needs only motif lists, it can be made for any pair of strains whose methylome is catalogued. Over 33,061,354 pairs of genomes of one species, formed by 39,920 genomes in REBASE [5], the fraction predicted separable from single reads rose with read length, from 36.97% at 1 kb to 74.56% at 10 kb and 91.21% at 50 kb (Fig. 3a). Two cautions apply. The database follows sequencing effort rather than biology, so *E. coli* alone contributes 51.15% of the pairs and four species account for 91.5% of them; and more than half of what is predicted separable at 1 kb rests on REBASE placeholder motifs. Every dataset in this study has median reads shorter than 10 kb, the left half of that curve (Fig. S3; Tables S34 to S36).

### Where methylation outperforms sequence-based tools, and where it does not

Two properties of a pair of strains are now known before any read is classified: their ANI, which bounds the sequence difference a read can carry, and their resolvability class, which bounds the methylation difference. Together they predict where Epilect should outperform tools that read sequence, and we tested that directly. We compared Epilect with the best of five configurations of three sequence-based read-phasing tools, Floria, devider and HairSplitter [7, 9, 10], choosing in each mixture whichever configuration scored highest (Methods). Choosing the best competitor after the fact favours the competitors. Two choices favour Epilect: it was given each strain’s methylome, built from that isolate’s reads not used in any mixture, which the sequence-based tools do not need; and its default settings were selected on this panel. Each of 176 pairings of two panel strains, one of them serving as the reference genome, was mixed computationally from each isolate’s own reads in four mixtures: 1:1, 1:4, 1:16, and 1:16 with the reference strain as the minority. Every tool received the same reads, and we scored each by per-read F1 over every read offered, so that a tool labelling few reads could not score as though it had answered the whole question (Tables S37, S73 and S78).

We defined a target regime of pairings from those two properties: an ANI of at least 99.50% together with a predicted methylome difference of any kind. Three of the six combinations this defines contain no pair anywhere in the panel, so the regime as tested consists of the three that do: pairs at 99.99% ANI or above with a coverage-only difference, and pairs between 99.50% and 99.99% with a coverage-only or a one-direction difference (Fig. 5a). Every pairing at 99.99% ANI or above is two isolates of one clone, so pairings of distinct strains occupy only the two combinations between 99.50% and 99.99%. The definition was drawn from the 12 pairings of the first benchmark that fall in the regime. Every other panel pairing meeting it, 62 in all (57 of distinct strains), was then selected on those two properties alone and run afterwards, alongside 62 pairings outside the regime matched on species and clone. These are new pairings but largely not new strains: they share 9 of the 10 clones that defined the regime, and the 7 pairings built only from other clones, 4 of them *L. monocytogenes*, are too few to test alone (0.08; 95% CI −0.06 to 0.28; Table S78). Within the target regime Epilect outperformed the best competitor. Over the 65 pairings of distinct strains, its per-read F1 exceeded the best competitor’s by 0.25 (95% CI 0.08 to 0.37; 260 mixtures, 18 clones), and Epilect scored higher in 228 of the 260 mixtures. Median per-read F1 was 0.47 for Epilect against 0.11 for the best competitor; we give these medians without an interval or a ratio, because the competitor’s median is close to zero and their ratio is unstable. Running every tool, Epilect included, at its own published defaults gave a margin of 0.21 (0.05 to 0.29). The advantage is one of accuracy rather than of answering where others cannot, since the best competitor returned output in every mixture and scored above zero in all 260. Including the nine pairings of two isolates from one clone, which enter the regime because their methylomes differ, the margin was 0.27 (0.08 to 0.38; 74 pairings, 296 mixtures) (Fig. 5b, c; Tables S73 and S78).

The advantage tracked the predicted methylome difference rather than the ANI range. In the 19 pairings of distinct strains from the same ANI range whose strains are predicted to share a methylome, Epilect scored lower than the best competitor, −0.18 (95% CI −0.19 to −0.13; 76 mixtures, 8 clones), so without a methylome difference the advantage reverses. Outside the target regime the margin was −0.05 (−0.21 to 0.12; 72 pairings of distinct strains), and there Epilect’s precision was lower than the best competitor’s, by 0.20 (0.06 to 0.35). Inside the regime the gain lay in recall: Epilect assigned more reads correctly, 0.20 (0.08 to 0.31), while its precision and the share of reads it labelled did not differ measurably from the best competitor’s (Fig. 5c; Table S78).

Three bounds apply to this result. First, it rests on one combination: between 99.50% and 99.99% ANI with a coverage-only difference the margin was 0.23 (95% CI 0.12 to 0.29; 46 pairings); with a one-direction difference the point estimate was larger but its interval crossed zero; and the seven pairings at 99.99% ANI or above, all of two isolates from one clone, came from only two clones, so no interval is given. Second, the advantage was detected only in skewed mixtures: at 1:1 the margin crossed zero (0.05, −0.09 to 0.16), whereas at the three skewed ratios it ran from 0.31 to 0.34, every interval excluding zero. Third, the point estimate for *L. monocytogenes* went the other way, on too few clones for an interval, and excluding it the margin was 0.27 (0.16 to 0.37). The result does not rest on *E. faecium* either, which contributes most target mixtures: excluding it left 0.18 (0.01 to 0.25) (Fig. 5c; Table S78).

No single level of sequence divergence divides the two approaches. Among pairings with a predicted methylome difference, and binned by the expected number of distinguishing single-nucleotide differences per read, the margin favoured Epilect below 5 differences per read, crossed zero between 5 and 25, and favoured the competitors at 25 or more (Table S78). Over all 704 mixtures the margin crossed zero (0.09, 95% CI −0.07 to 0.22; 0.09, −0.08 to 0.21 over the 548 mixtures of distinct strains). That figure describes the benchmark rather than bacteria, because 47% of the benchmark’s mixtures of distinct strains lie in the target regime (260 of 548), whereas the regime contains 65 of the 297 pairs above 99% ANI (21.9%; 95% CI 5.6 to 39.0; Table S78). The panel is not a random sample of strains, so neither share estimates how common such pairs are in nature. Membership of the target regime can be determined before sequencing from a pair’s ANI and motif lists, but outside it the data do not show where one approach gives way to the other.

### Assigning a locus to a strain

Laboratories usually act on a locus rather than a read: which of two co-resident strains carries a resistance gene or a plasmid-borne element. Having established where single reads can be assigned, we asked how many reads decide a locus. Requiring the locus score to reach 10, the fraction of cases in which ten covering reads identified the carrying strain correctly with probability at least 0.95 was 1.000 where both directions were predicted separable, 0.911 in one direction, 0.301 with coverage only and 0.000 where methylomes are identical, each case being one pairing with one true strain (Fig. 4b). Ten reads therefore decide a locus in the classes the prediction calls separable, and the prediction names those classes before any read is classified (Tables S47 and S48).

**Fig. 4.**
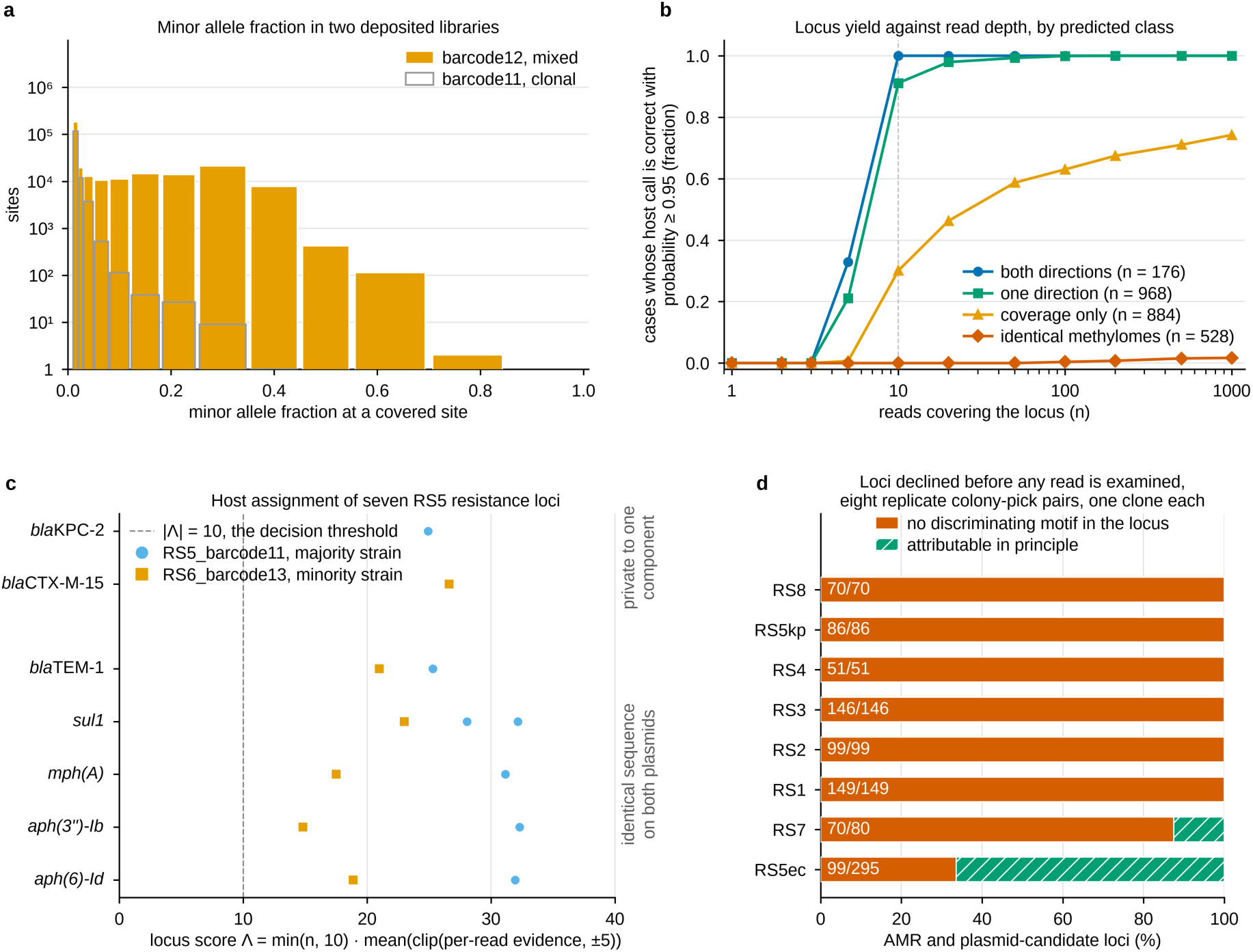
Assigning a locus to one of two co-resident strains. (a) Minor allele fraction at sites of depth at least 20 carrying a minor allele fraction of at least 0.01, in a two-strain deposited library and its clonal partner. (b) Fraction of cases in which the carrying strain is identified correctly with probability at least 0.95, against the number of covering reads, by predicted class, at default settings with a locus threshold of 10; a case is one pairing (two strains, one serving as the reference) with one true strain; *B* = 400 bootstrap draws, seed 20260913. (c) Locus score Λ for seven selected resistance loci of the two-strain library, five of them present at identical sequence on both strains’ plasmids. (d) Loci declined before any read is examined. Each of the eight PRJEB108076 pairs is two replicate colony picks of one clone, so their refusal rate is a specificity check; the rate on distinct strains is given in the text. See also Fig. S5.

The cap on the locus score, rather than the evidence itself, is what bounds false calls. We built two negative controls: one in which only one strain is present, so that every confident call is false, and one built from a motif both strains methylate. An uncapped sum of per-read evidence rose without limit, to false confident calls in 0.631 and 0.864 of cases at a thousand covering reads, whereas the capped score converged to 0.127 and 0.146. Where a control model could be built from a shared motif, false calls between isolates of one clone fell to zero, against 0.4286 where none could (Fig. S5; Table S50). A cross-run control using a single discriminating motif per strain gave spurious minority-strain fractions of 0.015 (*S. aureus* SA62) to 0.189 (*L. monocytogenes* LM41), tracking the motif’s methylation level (0.89 to 0.53) rather than the run or species; because it uses one motif, it does not bound the error of the classifier, which uses all of a strain’s motifs (Table S51).

A deposited isolate library provided a test with external ground truth. Library RS5_barcode12 holds a second strain at a median allele fraction of 0.376 over 124,556 diagnostic sites, beside its clonal partner in the same run (Fig. 4a; Table S52). Of 251 loci scored in that library, 170 were assigned, 61 were ambiguous and 20 were declined for want of signal, and all 112 loci that carried both decisive sequence provenance and an assignment were assigned to the correct strain. Five of those loci are present at identical sequence on both strains’ plasmids, where sequence alone cannot decide the host, and each copy was still assigned correctly (Fig. 4c and Fig. S6; Tables S52 to S54). How two strains came to share one library is undetermined, and we name no mechanism.

Epilect declines, before examining any read, a locus whose sequence contains no motif that distinguishes the two strains. Among the 297 pairs above 99% ANI it declined 11,233 of 19,679 resistance-gene and candidate plasmid loci (57.1%; 95% CI 43.1 to 70.3; Table S79). A third of this follows by arithmetic, from the 60 pairs that share a methylome entirely and therefore decline every locus. In the remaining 237 pairs, short loci were declined more often than long ones (64.6% of loci shorter than the median of 1,182 bp against 29.4% of longer loci; difference 35.2 percentage points, leave-one-clone-out 95% CI 21.7 to 48.6), as a longer locus has more chances to carry a distinguishing motif. Across all 297 pairs, resistance-gene loci were declined no more often than random genomic intervals of matched length (difference 3.1 percentage points, leave-one-clone-out 95% CI −1.7 to 7.8; Table S79). As expected, on eight pairs of replicate colony picks from a public collection, each pair two picks of a single clone, which carry no fixed sequence difference and in six of which no distinguishing motif, Epilect declined 770 of 976 loci (Fig. 4d; Table S55).

**Fig. 5.**
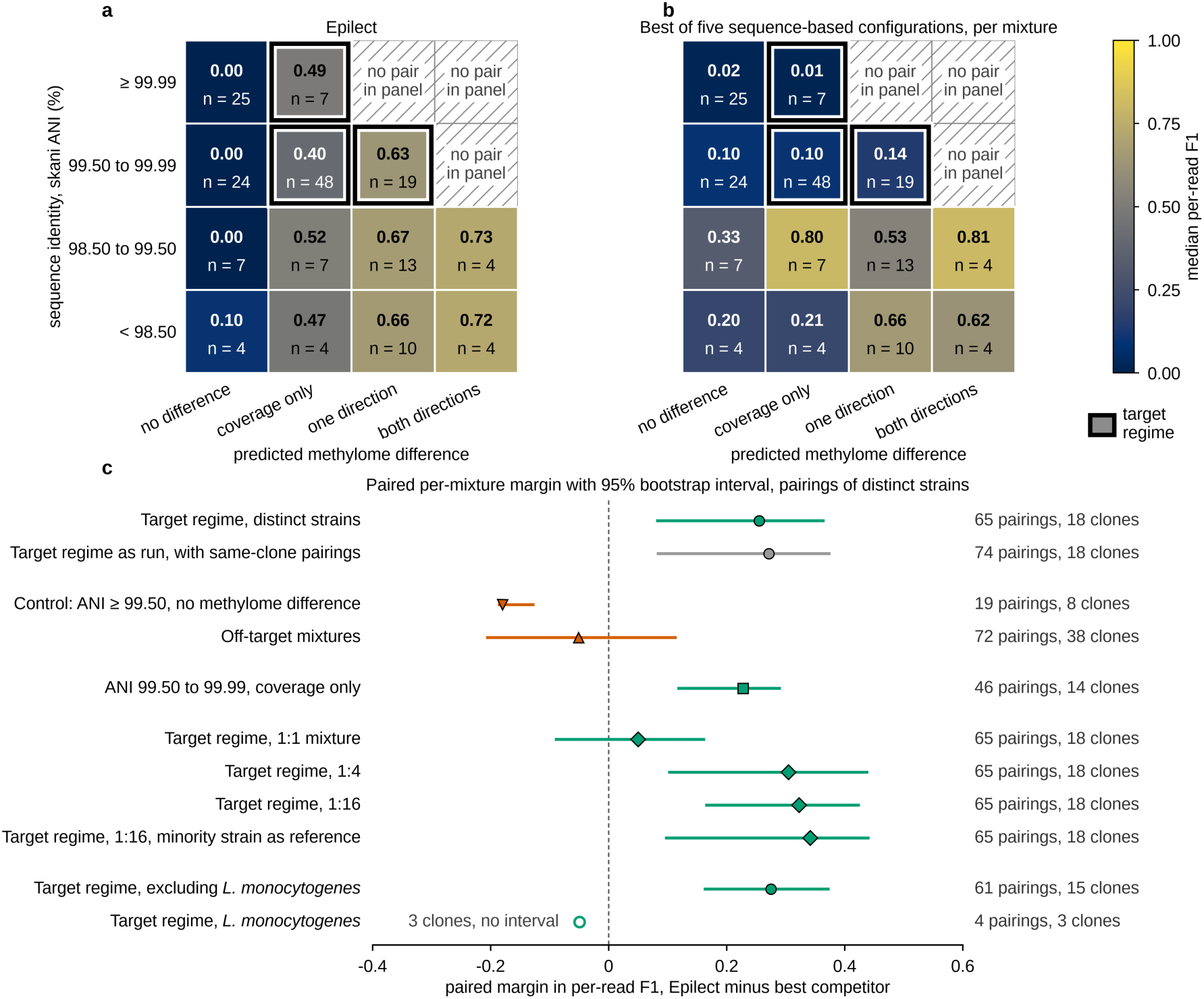
Epilect’s accuracy follows the predicted methylome difference. One benchmark of 704 computational mixtures: 176 pairings of two panel isolates, one serving as the reference, each mixed at 1:1, 1:4 and 1:16, the last in both orientations. Pairings are placed by ANI band (rows) and predicted methylome difference (columns); hatched combinations hold no pair in the panel, and the three occupied target combinations are outlined. (a) Median per-read F1 of Epilect over all 176 pairings. (b) Median per-read F1 of the best of five sequence-based configurations in each mixture. (c) Paired margin, Epilect minus the best competitor, on pairings of distinct strains: the target regime (65 pairings, 18 clones), the same regime as run including the nine pairings within one clone (74 pairings), the same-ANI control without a predicted methylome difference, the off-target regime, the coverage-only combination at 99.50% to 99.99% ANI, each mixing ratio, the target regime without *L. monocytogenes*, and *L. monocytogenes* alone. Bars are 95% dyadic cluster bootstrap intervals over clones (2,000 replicates); the open marker is a row with fewer than six clones. Values are in Table S78. See also Fig. S7.

## Discussion

Among isolates of one bacterial species above 99% ANI, 40.37% of read placements carried no sequence difference between the two strains, and the fraction rose to about three quarters above 99.9%. The rise held at every step of the pooled ladder and at eight of the twelve steps the four species could form. Any method that reads only sequence is bounded by this fraction, whatever its algorithm: it is the quantitative counterpart of the observation that one long-read strain-identification method grouped strains separated by only a few mutations [11], and it applies to the co-colonising lineages of *E. faecium*, *K. pneumoniae* and *S. aureus* that within-patient and transmission studies seek to separate [6, 12].

Methylation supplies a second source of evidence on those reads, within limits. Its precision on sequence-blind reads did not fall as strains converged, although its recall fell in *S. aureus*. In strains at 99.5% ANI or more that were predicted to differ in methylation, and in computational mixtures dominated by one strain, Epilect’s per-read F1 exceeded that of the best sequence-based tool. In equally related strains without a predicted methylome difference the comparison reversed, which is what would be expected if the advantage comes from methylation rather than from the pairs selected. On this evidence, Epilect and sequence-based phasing tools such as Floria, devider and HairSplitter [7, 9, 10] cover largely different pairs of strains, and the two are best used together. Methylation-aware tools such as Nanomotif and MicrobeMod [29, 30] work at the level of genomes and contigs; SMRT reads have been clustered by methylation without a reference methylome [31]; and CUPID links nanopore reads and contigs carrying resistance genes to their host species by methylation similarity [3]. Epilect addresses the step below the species: it assigns reads among strains of one species using a methylation profile supplied for each strain, and reports how often it is wrong.

That prediction of resolvability is the part of this work most likely to be useful to others. It needs only two strains’ motif lists, which can come from REBASE or from isolate sequencing, and across 639 pairs it placed 83.4% correctly among separation from single reads, by coverage only or not at all, erring more often towards caution than towards optimism. It did not anticipate which pairs would separate in both directions rather than one, and it was least reliable in *K. pneumoniae*. Because nanopore methylome profiles of an isolate have been reproducible between libraries [4] and strains of one species, even from one starter culture, frequently differ in their methylomes [23], a laboratory can estimate before sequencing whether a given pair of strains can be separated by methylation at its expected read length, and whether the pair falls in the regime where Epilect outperformed sequence-based tools. Settings in which the strains’ isolates are already sequenced and the question concerns their mixture fit this description: co-colonisation and relapse in a patient, an outbreak strain against a closely related background strain, the host of a plasmid shared between two co-resident strains, or a probiotic or starter strain tracked against a related resident strain.

Substitutions between these strains were clustered in all four species. In *E. faecium* we attribute that clustering to recombination by three criteria fixed in advance, although the signal rests on about three inferred ancestral events and the claim is made for one species only; in the other three species we name no mechanism. The resolvability class, not ANI, predicts whether methylation can separate two strains, because methylome difference is not a simple function of sequence difference.

### Limitations

The advantage over sequence-based tools is narrow. It rests on one combination of ANI and resolvability class, strains between 99.50% and 99.99% ANI with a coverage-only methylome difference; it was detected only in skewed mixtures and not at 1:1; its point estimate reversed in *L. monocytogenes*, on too few clones to estimate; and it depends on default settings selected on this panel. The two sources of evidence are positively correlated, so methylation is weakest where it is the only evidence, and its role narrows as reads lengthen: the sequence-blind fraction falls from 33.48% of read placements below 1 kb to 4.55% above 16 kb.

The prediction of resolvability was tested on the panel on which the default settings were chosen, with each isolate classified by its own methylation model and with motif lists discovered from the same reads, so its agreement with read assignment here is not an estimate for new strains. Its four-class agreement depended on the threshold that defines separability, and it was weakest in *K. pneumoniae*. The released tool implements a more conservative form of the rule than the one tested.

Every result on mixtures assumes a methylome supplied for each strain. Discovering the minority strain’s motifs from a mixture mapped to a single reference failed at every ratio tested, including 1:1, so without a reference methylome or an assembly that separates the strains no two-strain model can be built. Methylomes are also not fixed: methylation can vary within a clonal population through phase-variable methyltransferases [52], and masking phase-variable loci did not improve the results here (Methods). Between two sequencing runs of the same *S. aureus* or *E. faecium* isolate, 0.18% to 7.37% of loci differed in methylation by at least 20 percentage points, against 4.93% between two isolates of one clone and 47.50% and 76.52% between strains at 99.02% and 99.44% ANI (loci with at least 10 reads; Table S49). We expect that a methylome measured on one isolate may therefore not describe every cell of the strain, or the same strain sampled later.

The default settings favour precision over recall. The fidelity filter supplies essentially all of the precision gain and costs recall on sequence-blind read placements, while the lower threshold with a one-site minimum supplies the recall. About one read placement in seven never reaches the classifier, 13.45% carrying no methylation call at all and the fidelity filter emptying a further 1.02 percentage points. Locus attribution declines 57.1% of loci on pairs above 99% ANI, a third of that because 60 pairs share a methylome entirely. At default settings a sample needs several gigabytes of memory, which can be reduced below a quarter of a gigabyte at the cost of wall time.

The panel comprises 79 isolates but 50 clones, and one species supplies 137 of the 297 pairs above 99% ANI from 8 clones, so pooled figures reflect its composition: removing its most influential clone moves the sequence-blind fraction from 40.37% to 35.19%. The fraction also depends on the definition of sequence-blindness, the alignment setting and the ANI threshold (Methods). The panel cannot settle whether methylome divergence is unrelated to sequence divergence: the two are associated in *S. aureus*, the pooled estimate straddles zero, and an equivalence test is underpowered in either direction. Finally, the evidence comes from isolate panels, computational mixtures, one two-strain library and one patient specimen, not from clinical or environmental samples with isolate ground truth. The next test is a set of such samples in which the co-resident strains have been isolated and their methylomes measured.

Reads from closely related strains often carry no sequence evidence of their origin. Methylation recovers part of that evidence with high precision for pairs of strains whose methylomes differ, and those pairs can largely be identified from motif lists before any sample is sequenced.

## Author statements

### Author contributions

Kardokh Kakabra: conceptualization, methodology, software, validation, formal analysis, investigation, data curation, visualization, writing (original draft), writing (review and editing). Marcus J. Claesson: conceptualization, resources, supervision, project administration, funding acquisition, writing (review and editing).

### Ethical approval

Not applicable. This study reanalysed publicly deposited bacterial sequencing data and generated no new data from humans or animals.

## Acknowledgements

The authors wish to acknowledge the Irish Centre for High-End Computing (ICHEC) for the provision of computational facilities and support through the Interim National HPC Service on the MeluXina supercomputer operated by LuxProvide.

## Use of generative AI

During the preparation of this work the authors used Claude (Anthropic; model Claude Opus 5.5) to assist with the editorial process. Specifically, the model was employed to refine the academic language, enhance the overall clarity and flow of the text, and ensure consistency in terminology. After using this tool, the authors reviewed and edited the content as needed and take full responsibility for the content of the published article.

## Supplementary material

This file contains Figs S1 to S9, each with its legend. The supplementary tables are provided separately as the Excel workbook Supplementary_Tables_S1-S80.xlsx, with a Contents sheet listing each table’s number, title and description.

**Fig. S1.**
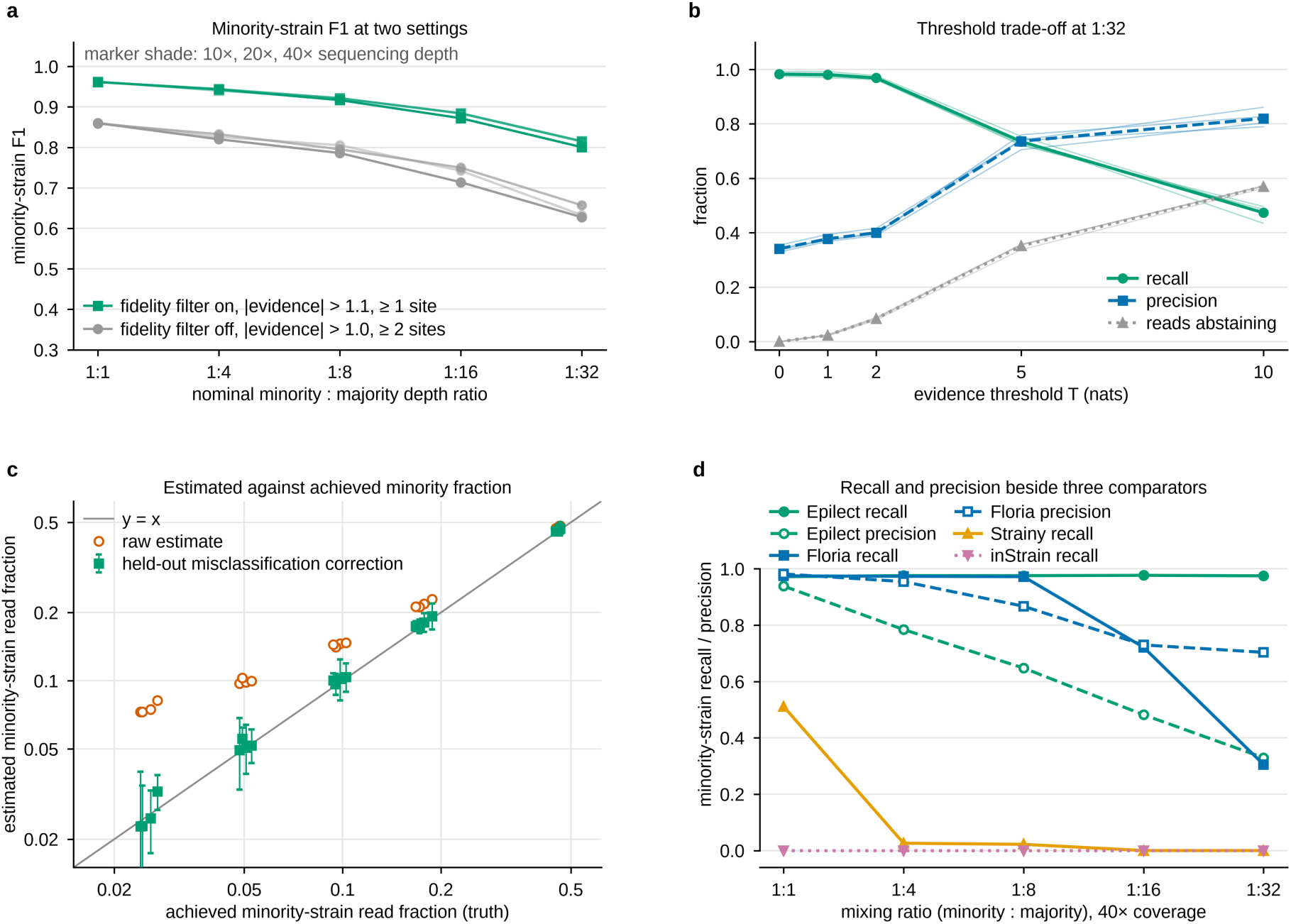
Minority-strain read assignment on the end-to-end mock of two *Streptococcus dysgalactiae* subsp. *equisimilis* (SDSE) isolates. (a) Minority-strain F1 (harmonic mean of precision and recall) at five minority : majority depth ratios and three depths (15 mixtures), at default settings (fidelity filter on, |evidence| > 1.1, at least one informative site) and with the filter off (|evidence| > 1.0, two sites). (b) Recall, precision and the fraction of reads abstaining against the evidence threshold T at 1:32; pale lines are four coverages (5× to 40×), heavy lines their mean. At 40×, raising T from 0 to 5 raised precision from 0.330 to 0.705 and lowered recall from 0.975 to 0.721, with 35.8% of reads abstaining. (c) Estimated against achieved minority read fraction at T = 0, before and after held-out misclassification correction, in the held-out half of 20 mixtures; error bars are sampling intervals on the corrected estimate (0.94 to 1.20 of the achieved fraction). (d) Minority-strain recall and precision at 40× beside Floria, Strainy and inStrain; recall denominators differ (Epilect, 560 true-minority reads with a GATC or CCWGG call; Floria, 396 reads placed in a haploset; 695 offered). The Strainy series comes from a separate baseline run, so its precision cannot be recomputed, and inStrain assigned no reads, so its precision is undefined. Panels b to d were measured with the filter off and a one-site floor.

**Fig. S2.**
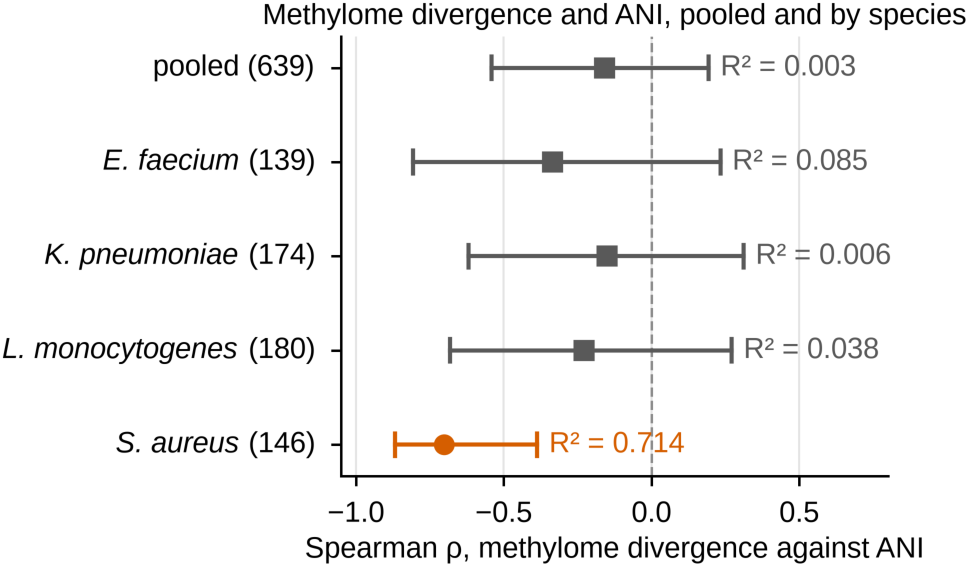
Methylome divergence against average nucleotide identity (ANI), pooled and by species. Spearman ρ between the number of motifs whose methylated fraction differs by at least 0.5 and pairwise skani ANI, pooled over 639 genuine conspecific pairs and within each species (number of pairs in parentheses: *Enterococcus faecium* 139, *Klebsiella pneumoniae* 174, *Listeria monocytogenes* 180, *Staphylococcus aureus* 146). Bars are 95% dyadic cluster bootstrap intervals (2,000 replicates) over 50 clones, a clone being a single-linkage cluster of isolates at ANI 99.99%; R² from the corresponding Pearson fit is printed beside each. Pooled, ρ = −0.159 (95% CI −0.541 to 0.194; R² = 0.003). In *S. aureus* ρ = −0.702 (−0.868 to −0.387; R² = 0.714), the only interval excluding zero. The 139 *E. faecium* pairs come from 8 clones. Nineteen of the 639 pairs involve isolate Kp17, a *Klebsiella quasipneumoniae* subsp. *similipneumoniae*; removing them moves the *K. pneumoniae* coefficient from −0.151 to −0.167 (−0.649 to 0.360). An interval that includes zero is not evidence of no association: among the 297 pairs above 99% ANI, pooled ρ was −0.293 (−0.648 to 0.056) and *S. aureus* ρ −0.820 (−0.979 to −0.378; 34 pairs, 12 clones).

**Fig. S3.**
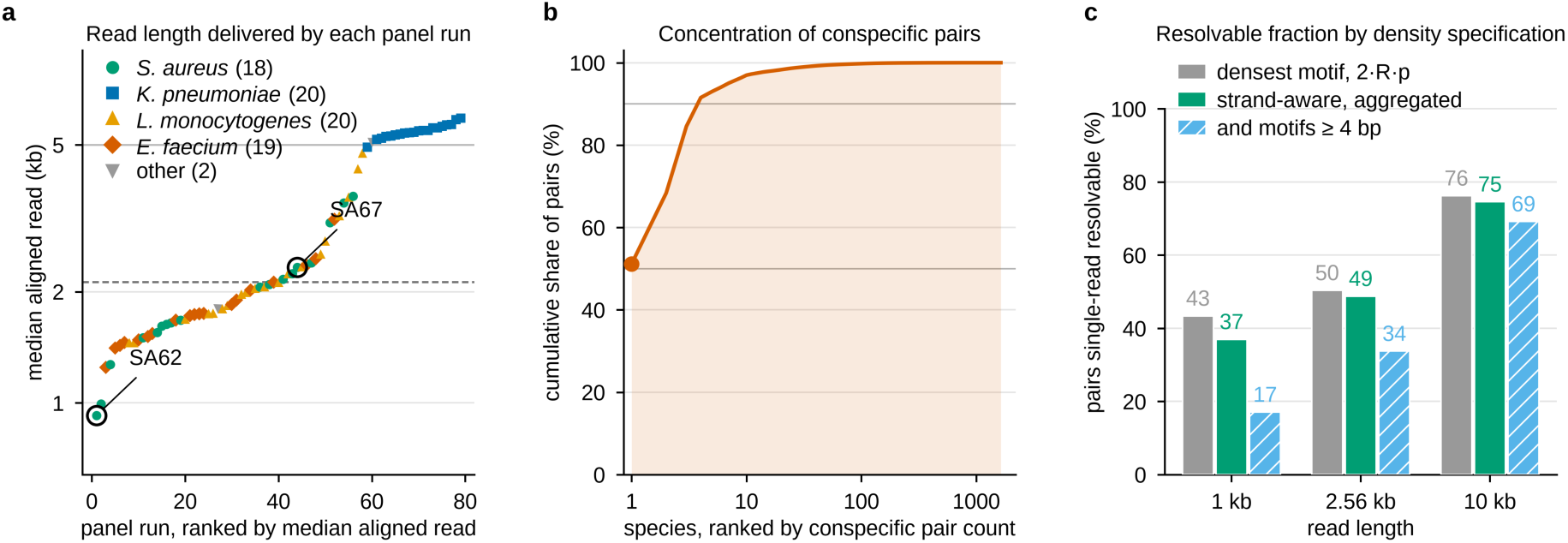
Read lengths in this study, and how the resolvability prediction depends on its density specification. (a) Median aligned read length of each of the 79 panel runs, ranked and coloured by species, over 11,165,884 aligned reads. The dashed rule is the panel median, 2,124 bp; the solid rule is 5 kb, reached by 20 of 79 runs. The two *Staphylococcus aureus* isolates SA62 and SA67 are circled: SA62 is the shortest run (925 bp median) and SA67 ranks 44th (2,327 bp). (b) Cumulative share of the 33,061,354 conspecific pairs in the REBASE map against species rank, with grey rules at 50% and 90%; *Escherichia coli* contributes 51.1% (16,910,020 pairs) and the top four species 91.5% (30,258,530), and 1,637 species contribute at least two strains. (c) Percentage of pairs predicted single-read resolvable at three read lengths under three density specifications: the densest-motif rule 2·R·p (expected discriminating sites per read from the single densest private motif, twice the read length R times the motif’s per-base occurrence probability p, counting both strands), the strand-aware aggregated rule used throughout, and that rule restricted to motifs of at least 4 bp, which removes REBASE placeholder motifs. At 1 kb these gave 43.4% (14,336,726 of 33,061,354 pairs), 37.0% (12,221,729 of 33,061,354) and 17.1% (of 32,932,469), and at 10 kb 76.2% (25,197,404), 74.6% (24,650,829) and 69.2% (of 32,932,469).

**Fig. S4.**
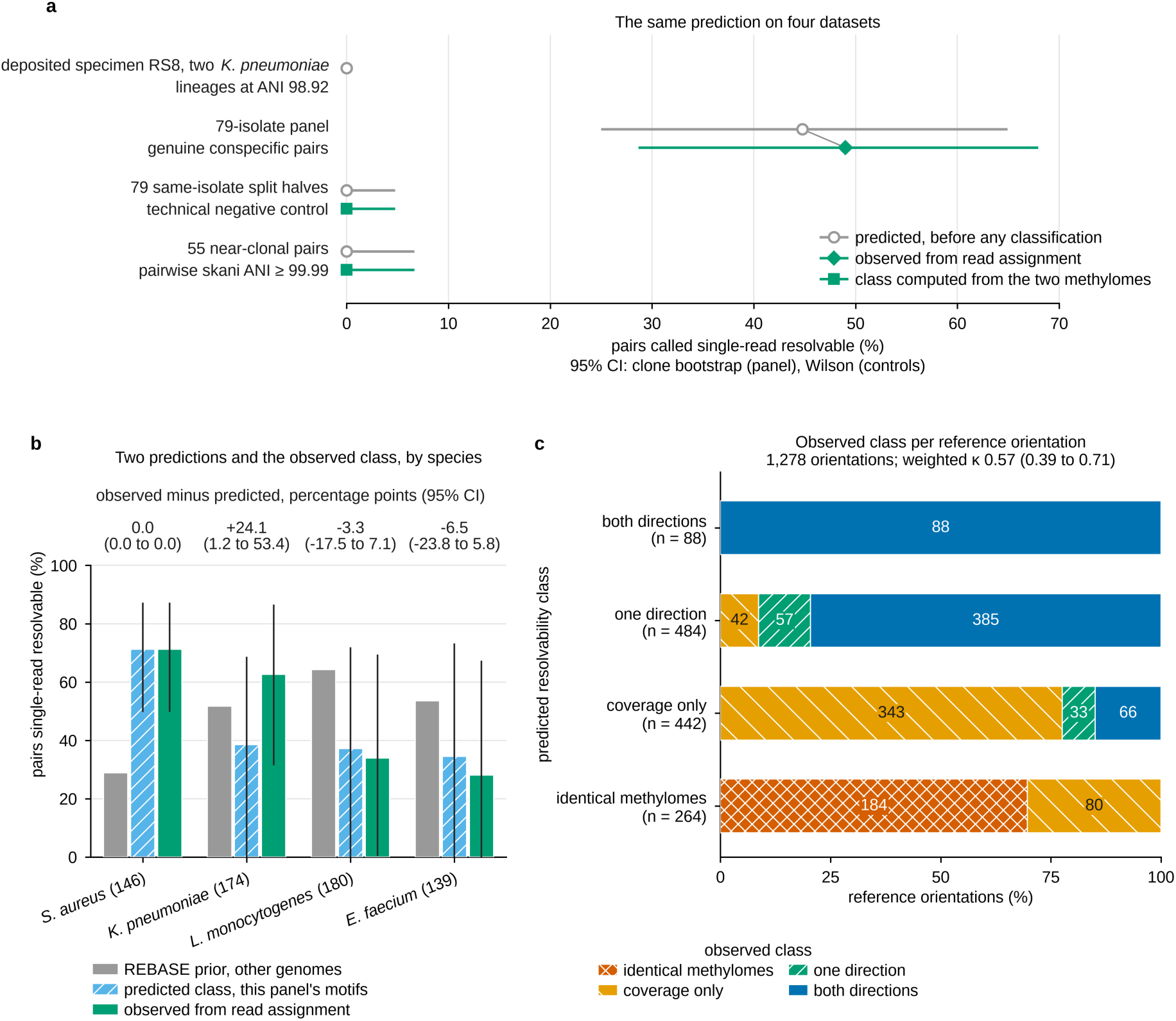
The resolvability prediction tested on four datasets, by species and per reference orientation. (a) Pairs predicted single-read resolvable (circles) against, for the panel, the fraction observed from read assignment at default settings (diamond) and, for the two controls, the class computed from the two methylomes (squares), which is not a read-assignment outcome. Top to bottom: specimen RS8, two *Klebsiella pneumoniae* lineages at 98.92% ANI, predicted not resolvable (n = 1, no per-read outcome); the 639 genuine conspecific pairs of the 79-isolate panel, 44.8% predicted (242 one direction, 44 both directions) and 49.0% (313) observed, the class agreeing in 333 (52.1%; linearly weighted kappa 0.57, 95% CI 0.39 to 0.71); 79 same-isolate split halves, 0 of 79; 55 near-clonal pairs (ANI ≥ 99.99%), 0 of 55. The panel’s intervals are clone bootstraps over 50 clones, because pairs share isolates; the controls’ are Wilson intervals that treat units as independent. (b) By species, a REBASE prior from unrelated genomes, the class predicted from this panel’s motifs and the observed class, with clone-bootstrap intervals (none on the prior). Observed minus predicted in percentage points (95% CI): *Staphylococcus aureus* 0.0 (0.0 to 0.0), *K. pneumoniae* +24.1 (1.2 to 53.4), *Listeria monocytogenes* −3.3 (−17.5 to 7.1) and *Enterococcus faecium* −6.5 (−23.8 to 5.8; 8 clones). (c) Observed class in each of the 1,278 reference orientations (each strain once the reference), by predicted class, under the same rule; the class agreed in 672 (52.6%; kappa 0.57, 0.39 to 0.71).

**Fig. S5.**
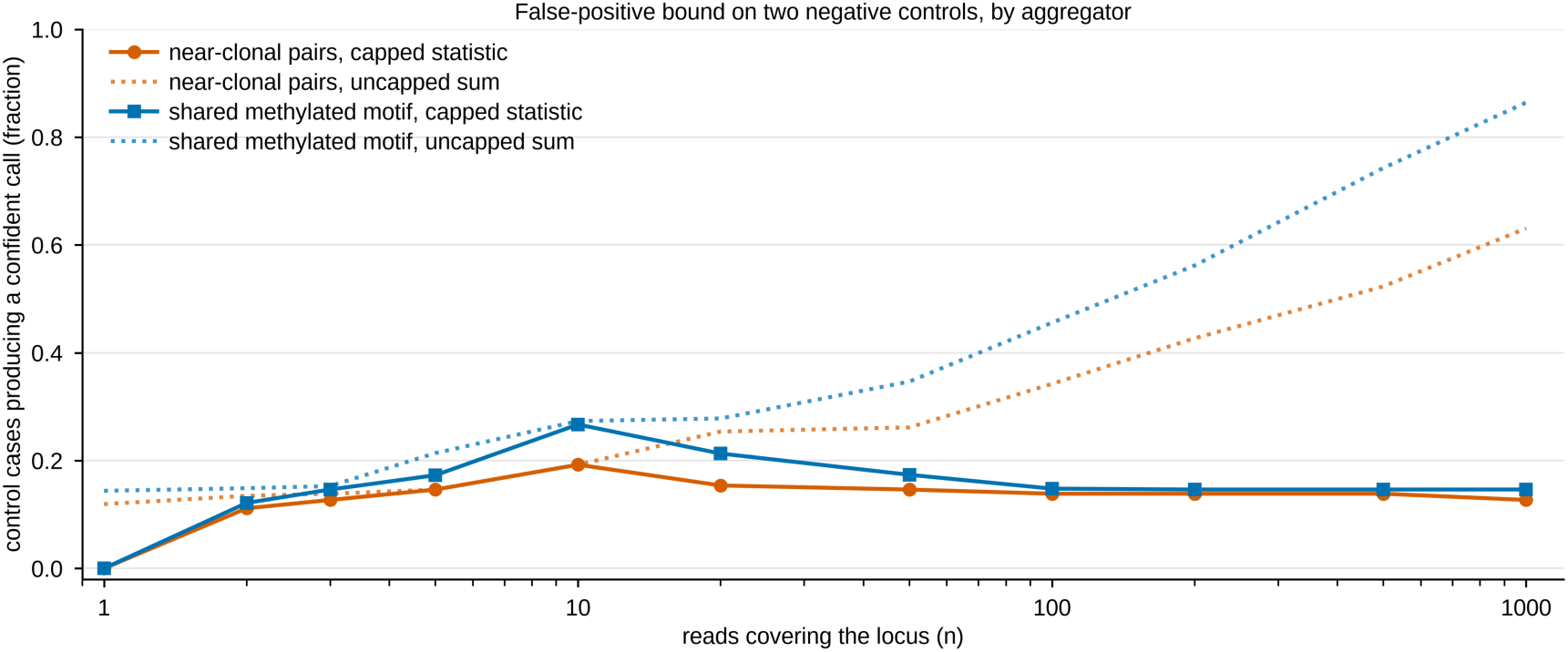
Capping the locus score bounds false confident calls on two negative controls. Fraction of control cases producing a confident host call more than 1% of the time, against the number of reads covering the locus, under the locus score Λ = min(n, 10) × mean per-read evidence clipped at ±5, where n is the number of reads covering the locus (capped, solid), and under an uncapped sum of per-read evidence (dotted). The controls are near-clonal pairs, where only one strain is present so every confident call is false (n = 260 cases), and a shared methylated motif, a model that cannot discriminate (n = 1,244 cases). At a thousand reads the uncapped sum reached 0.631 and 0.864; the capped score converged to 0.127 and 0.146. Near-clonal false positives concentrate where no shared-motif control model can be built: 0.4286 of cases (n = 56) against 0.0000 where one exists (n = 204), at a locus score threshold of 10 (source data only). Measured at default settings, with reads the fidelity filter empties of calls counted at zero evidence.

**Fig. S6.**
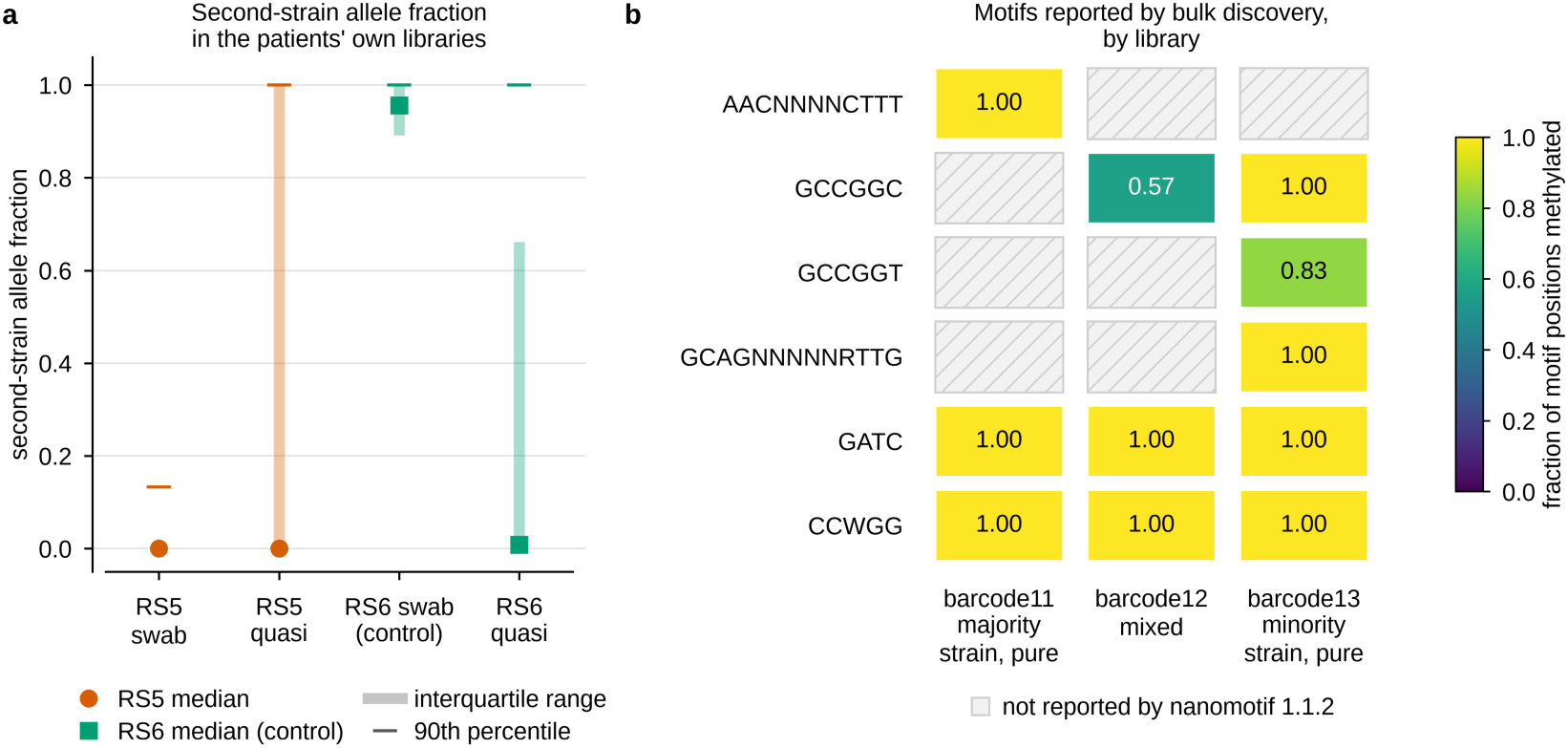
The two-strain deposited library in detail. (a) Second-strain allele fraction at diagnostic sites in the two patients’ own native metagenomes and quasi-metagenomes (the depositors’ name, in ENA PRJEB108076, for a second library prepared from the same rectal swab): median (circle for RS5, square for the RS6 positive control), interquartile range (bar) and 90th percentile (tick), over 78,719 to 123,444 sites per library at median depths of 12× to 124×. The second strain of library RS5_barcode12 (patient RS5’s deposited isolate library on barcode 12) is absent from patient RS5’s own swab (median 0.000, 90th percentile 0.133), whereas the positive control from patient RS6 gives a median of 0.957. (b) Fraction of motif positions methylated for six motifs in the majority strain’s pure library, the mixed library and the minority strain’s pure library, as reported by bulk motif discovery (nanomotif 1.1.2); hatched tiles are motifs not reported for that library. Bulk discovery on the mixed library fails in both directions: AACNNNNCTTT is reported for the pure majority library (1.000) but not for the mixture, and GCCGGC is reported for the mixture at 0.567 although it belongs to the minority strain (0.998 in its own library). The mechanism that placed two strains in one deposited library is undetermined.

**Fig. S7.**
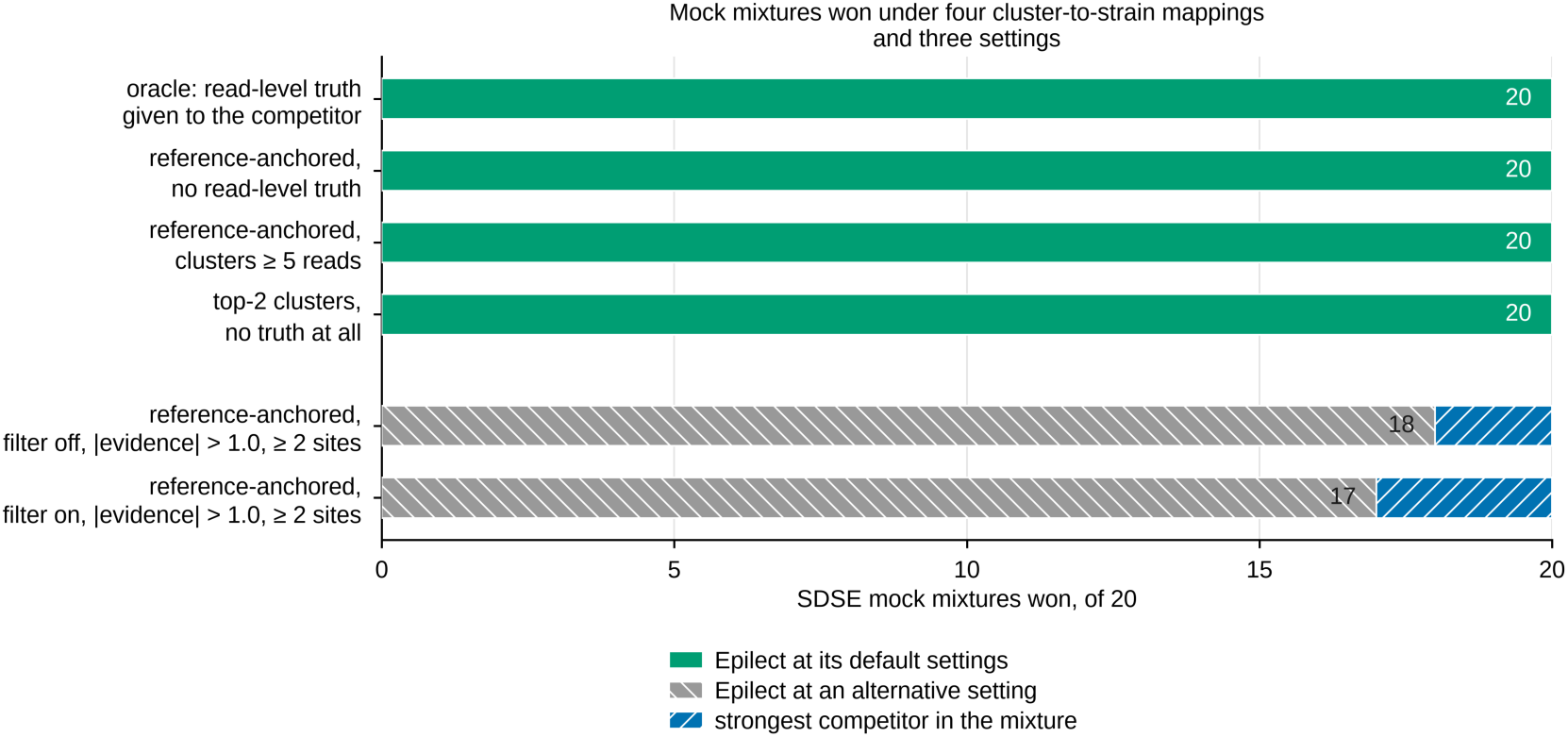
At default settings Epilect wins all 20 SDSE mock mixtures under four rules for mapping a competitor’s read clusters to strains. Number of the 20 mixtures of two *Streptococcus dysgalactiae* subsp. *equisimilis* (SDSE) isolates in which Epilect’s per-read F1 (harmonic mean of precision and recall) over every offered read exceeds that of the strongest competitor in the same mixture. That every tool was scored over one offered read set is asserted rather than counted: all 20 runs record an identical scored-read source, but no mixture was recounted read by read. Mappings differ in the truth the competitor receives: top-2 clusters, none; reference-anchored mapping labels a cluster by the strain-private alleles its consensus carries, with and without a five-read minimum cluster size; the oracle mapping supplies read-level truth. At default settings (fidelity filter on, |evidence| > 1.1, at least one informative site) Epilect won 20 of 20 mixtures under every mapping. Hatched bars give two alternative settings on the reference-anchored mapping: 18 of 20 with the filter off at |evidence| > 1.0 and at least two informative sites, and 17 of 20 with the filter on at the same threshold. Competitors were Floria, Floria -p 2, devider, devider --sensitive, HairSplitter and Strainy; Strainy is not reproducible run to run because metaFlye pins no seed. These mixtures overstate the margin relative to the 704-mixture benchmark of Fig. 5.

**Fig. S8.**
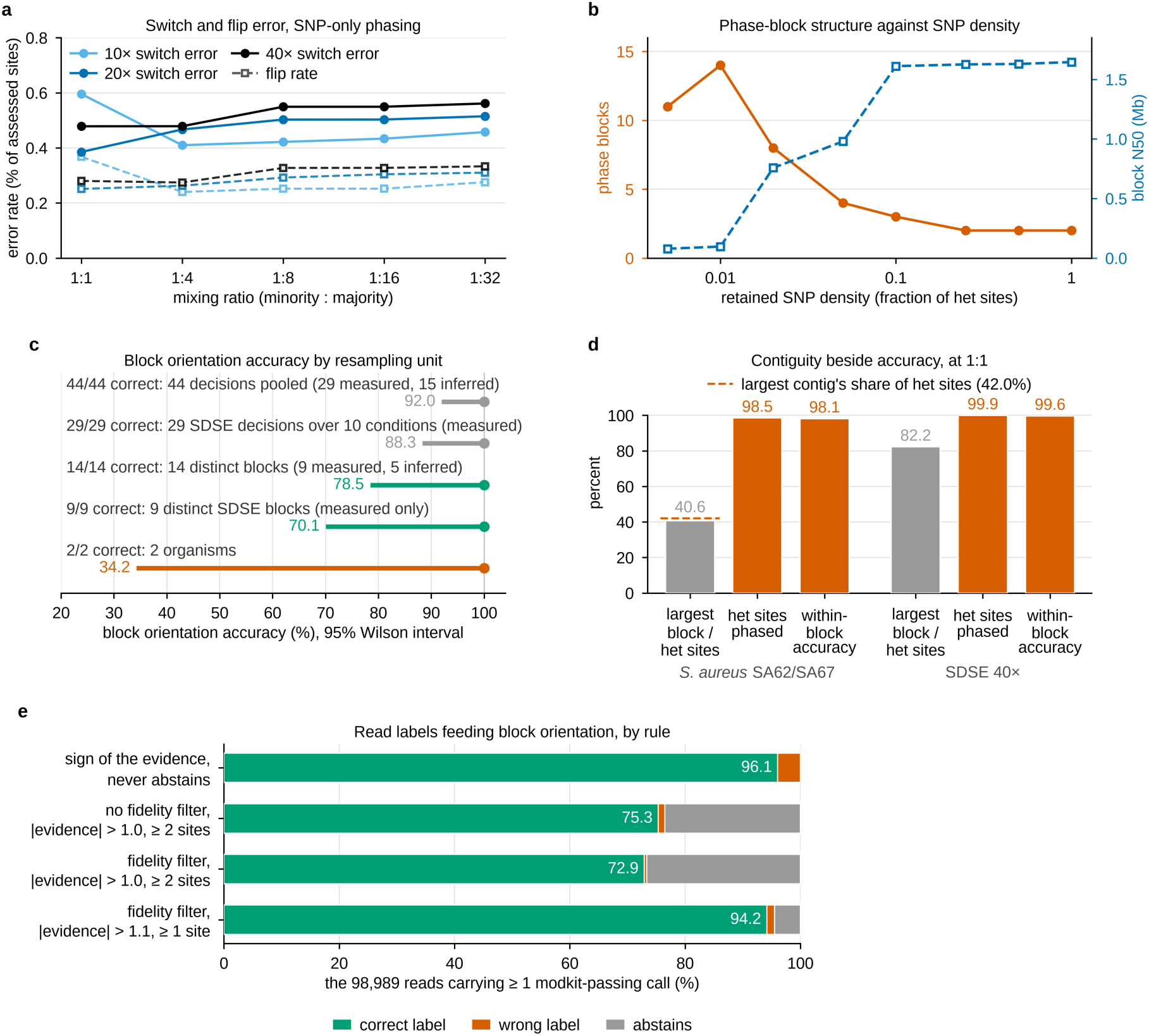
Single-nucleotide polymorphism (SNP)-based phasing of the mock mixtures, and the methylation read labels that orient its blocks. (a) Switch error (solid) and flip rate (dashed) for read-backed SNP-only phasing of the mock mixtures of two *Streptococcus dysgalactiae* subsp. *equisimilis* (SDSE) isolates (Fig. S1) at three depths and five mixing ratios; over the 15 mixtures switch error stayed between 0.39% and 0.60% of assessed sites. (b) Phase blocks (left axis) and block N50 (right axis; the length at which blocks that long or longer cover half the phased sequence) as heterozygous (het) sites are removed. (c) Block orientation accuracy with 95% Wilson intervals under five choices of independent unit (grey: decisions; green: distinct blocks; vermilion: organisms); every decision was correct, and the lower bound is 92.0% over 44 pooled decisions, 78.5% over 14 distinct blocks and 34.2% over two organisms. (d) At 1:1, the largest block as a percentage of het sites, het sites phased and within-block accuracy, for *Staphylococcus aureus* SA62/SA67 (29,885 het sites) and SDSE at 40× (17,154); the dashed line is the largest contig’s share of het sites (42.0%), which caps the first statistic. (e) Read labels under four rules, over the 98,989 pure control reads with at least one call passing modkit 0.6.4 filtering: the sign of the evidence labels 96.06% correctly; default settings label 94.18% correctly, with 1,330 errors and 4,434 abstentions.

**Fig. S9.**
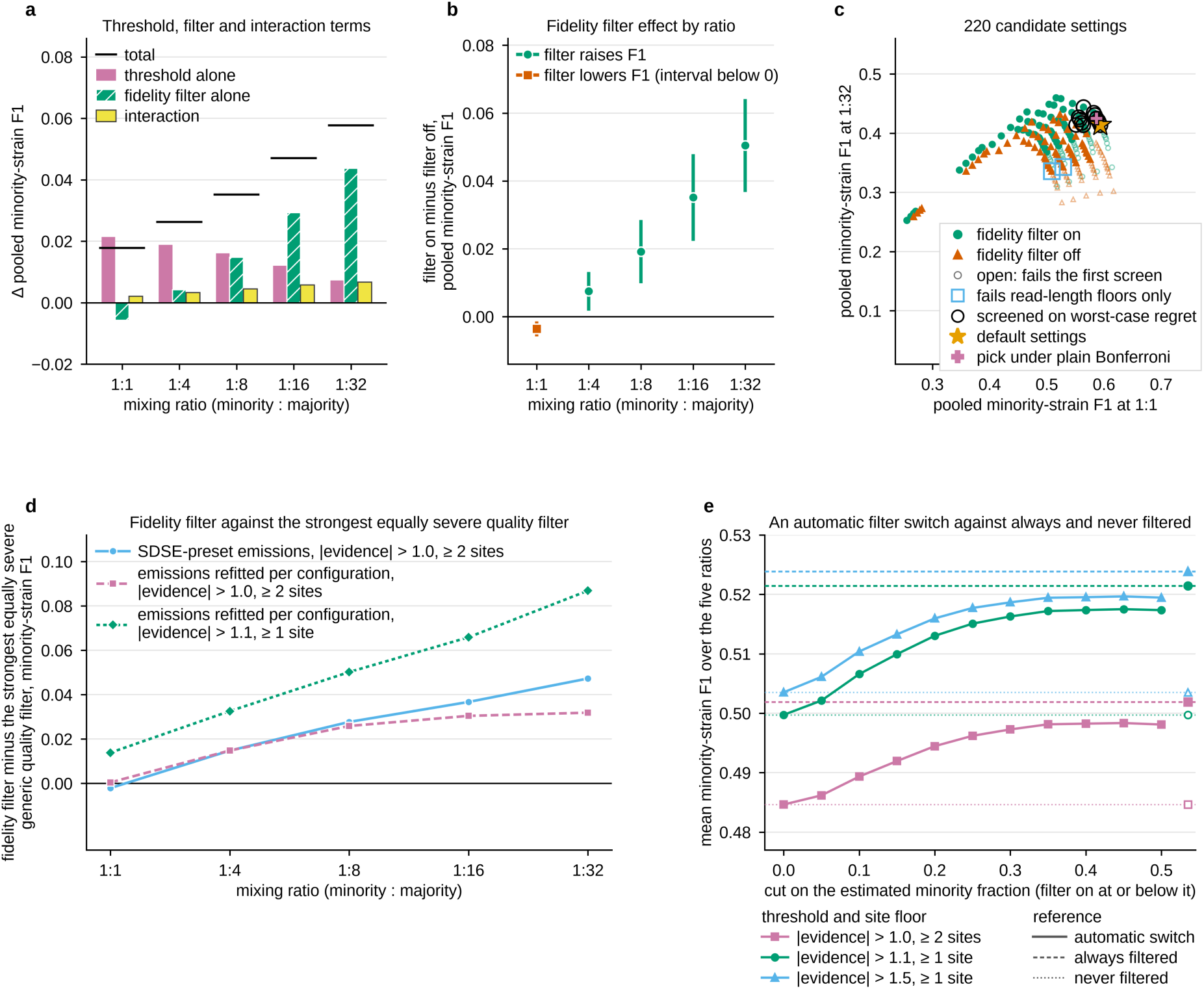
Selection of the default settings. Panels a to c give pooled minority-strain F1 (harmonic mean of precision and recall), and panel e its mean over the five ratios, over 1,278 genuine non-near-clonal pairings (two strains, one the reference genome), each scored with either isolate as the minority (2,556 mixtures); panel d is the SDSE mock (Fig. S1), three mixtures per ratio. (a) Change in pooled F1 from the filter off at |evidence| > 1.0 and two sites, split into threshold, filter and interaction terms; the black rule is the total. (b) Effect of the fidelity filter alone at the default threshold, with paired leave-one-clone-out 95% intervals over 50 clones (circles where the filter raises F1, the square where it lowers it). (c) All 220 candidate settings, by pooled F1 at 1:1 against 1:32; open markers fail the first screen. Twelve matched or beat the default’s worst-case regret and went to a second screen of 110 stratum-level tests per candidate (170 with six read-length bins, same pick; Table S56). Only the default passed under familywise control by the family’s maximum-|z| null (critical value 3.205 for the 110 tests), Holm, Bonferroni at the effective number of tests or Benjamini-Hochberg; under plain Bonferroni three points pass and the pick would move to the rival marked +. (d) The fidelity filter minus the strongest equally severe generic quality filter, in three configurations; each point is a mean of three mixtures, without interval. (e) An automatic filter switch set by the estimated minority fraction; dashed lines are always filtered and dotted lines never filtered.

