## Supplementary for "DNA methylation assigns sequencing reads to closely related bacterial strains when their sequence cannot": Supplementary_Material_2026-11-07.pdf


### Kardokh Kakabra and Marcus J. Claesson


This file contains Figs S1 to S9, each with its legend. The supplementary tables are provided separately as the Excel workbook Supplementary_Tables_S1-S80.xlsx, with a Contents sheet listing each table's number, title and description.

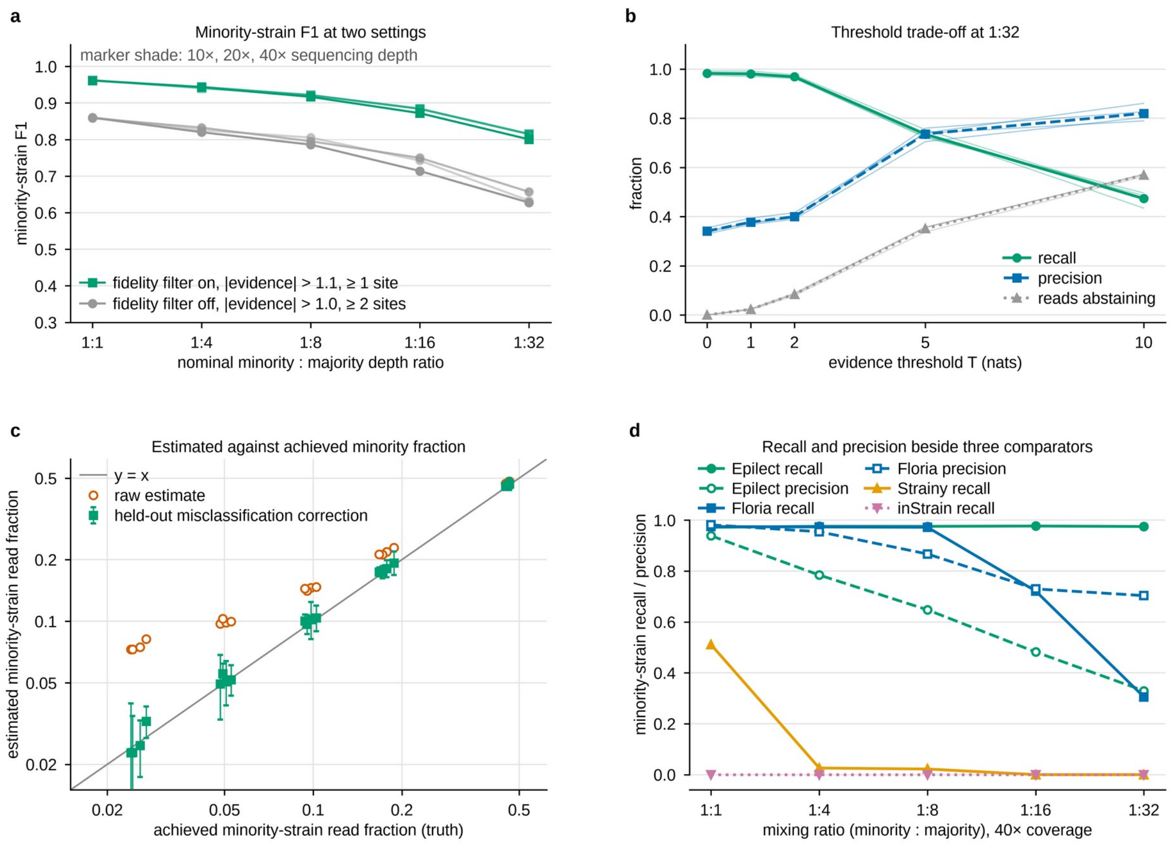


**Fig. S1.** Minority-strain read assignment on the end-to-end mock of two *Streptococcus dysgalactiae* subsp. *equisimilis* (SDSE) isolates. (a) Minority-strain F1 (harmonic mean of precision and recall) at five minority : majority depth ratios and three depths (15 mixtures), at default settings (fidelity filter on, |evidence| > 1.1, at least one informative site) and with the filter off (|evidence| > 1.0, two sites). (b) Recall, precision and the fraction of reads abstaining against the evidence threshold T at 1:32; pale lines are four coverages (5× to 40×), heavy lines their mean. At 40×, raising T from 0 to 5 raised precision from 0.330 to 0.705 and lowered recall from 0.975 to 0.721, with 35.8% of reads abstaining. (c) Estimated against achieved minority read fraction at T = 0, before and after held-out misclassification correction, in the held-out half of 20 mixtures; error bars are sampling intervals on the corrected estimate (0.94 to 1.20 of the achieved fraction). (d) Minority-strain recall and precision at 40× beside Floria, Strainy and inStrain; recall denominators differ (Epilect, 560 true-minority reads with a GATC or CCWGG call; Floria, 396 reads placed in a haploset; 695 offered). The Strainy series comes from a separate baseline run, so its precision cannot be recomputed, and inStrain assigned no reads, so its precision is undefined. Panels b to d were measured with the filter off and a one-site floor.

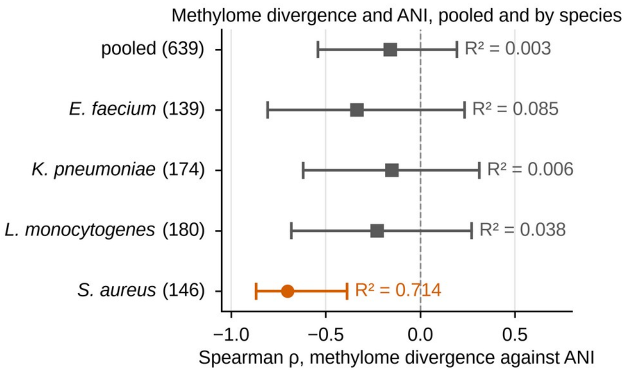


**Fig. S2.** Methylome divergence against average nucleotide identity (ANI), pooled and by species. Spearman ρ between the number of motifs whose methylated fraction differs by at least 0.5 and pairwise skani ANI, pooled over 639 genuine conspecific pairs and within each species (number of pairs in parentheses: *Enterococcus faecium* 139, *Klebsiella pneumoniae* 174, *Listeria monocytogenes* 180, *Staphylococcus aureus* 146). Bars are 95% dyadic cluster bootstrap intervals (2,000 replicates) over 50 clones, a clone being a single-linkage cluster of isolates at ANI 99.99%; $R^2$ from the corresponding Pearson fit is printed beside each. Pooled, ρ = −0.159 (95% CI −0.541 to 0.194; $R^2$ = 0.003). In *S. aureus* ρ = −0.702 (−0.868 to −0.387; $R^2$ = 0.714), the only interval excluding zero. The 139 *E. faecium* pairs come from 8 clones. Nineteen of the 639 pairs involve isolate Kp17, a *Klebsiella quasipneumoniae* subsp. *similipneumoniae*; removing them moves the *K. pneumoniae* coefficient from −0.151 to −0.167 (−0.649 to 0.360). An interval that includes zero is not evidence of no association: among the 297 pairs above 99% ANI, pooled ρ was −0.293 (−0.648 to 0.056) and *S. aureus* ρ −0.820 (−0.979 to −0.378; 34 pairs, 12 clones).

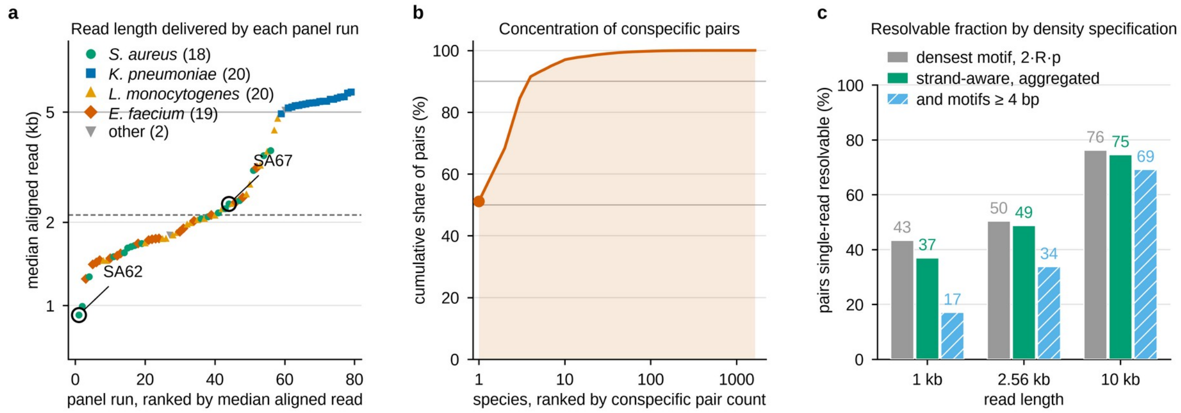


**Fig. S3.** Read lengths in this study, and how the resolvability prediction depends on its density specification. (a) Median aligned read length of each of the 79 panel runs, ranked and coloured by species, over 11,165,884 aligned reads. The dashed rule is the panel median, 2,124 bp; the solid rule is 5 kb, reached by 20 of 79 runs. The two *Staphylococcus aureus* isolates SA62 and SA67 are circled: SA62 is the shortest run (925 bp median) and SA67 ranks 44th (2,327 bp). (b) Cumulative share of the 33,061,354 conspecific pairs in the REBASE map against species rank, with grey rules at 50% and 90%; *Escherichia coli* contributes 51.1% (16,910,020 pairs) and the top four species 91.5% (30,258,530), and 1,637 species contribute at least two strains. (c) Percentage of pairs predicted single-read resolvable at three read lengths under three density specifications: the densest-motif rule 2·R·p (expected discriminating sites per read from the single densest private motif, twice the read length R times the motif's per-base occurrence probability p, counting both strands), the strand-aware aggregated rule used throughout, and that rule restricted to motifs of at least 4 bp, which removes REBASE placeholder motifs. At 1 kb these gave 43.4% (14,336,726 of 33,061,354 pairs), 37.0% (12,221,729 of 33,061,354) and 17.1% (of 32,932,469), and at 10 kb 76.2% (25,197,404), 74.6% (24,650,829) and 69.2% (of 32,932,469).

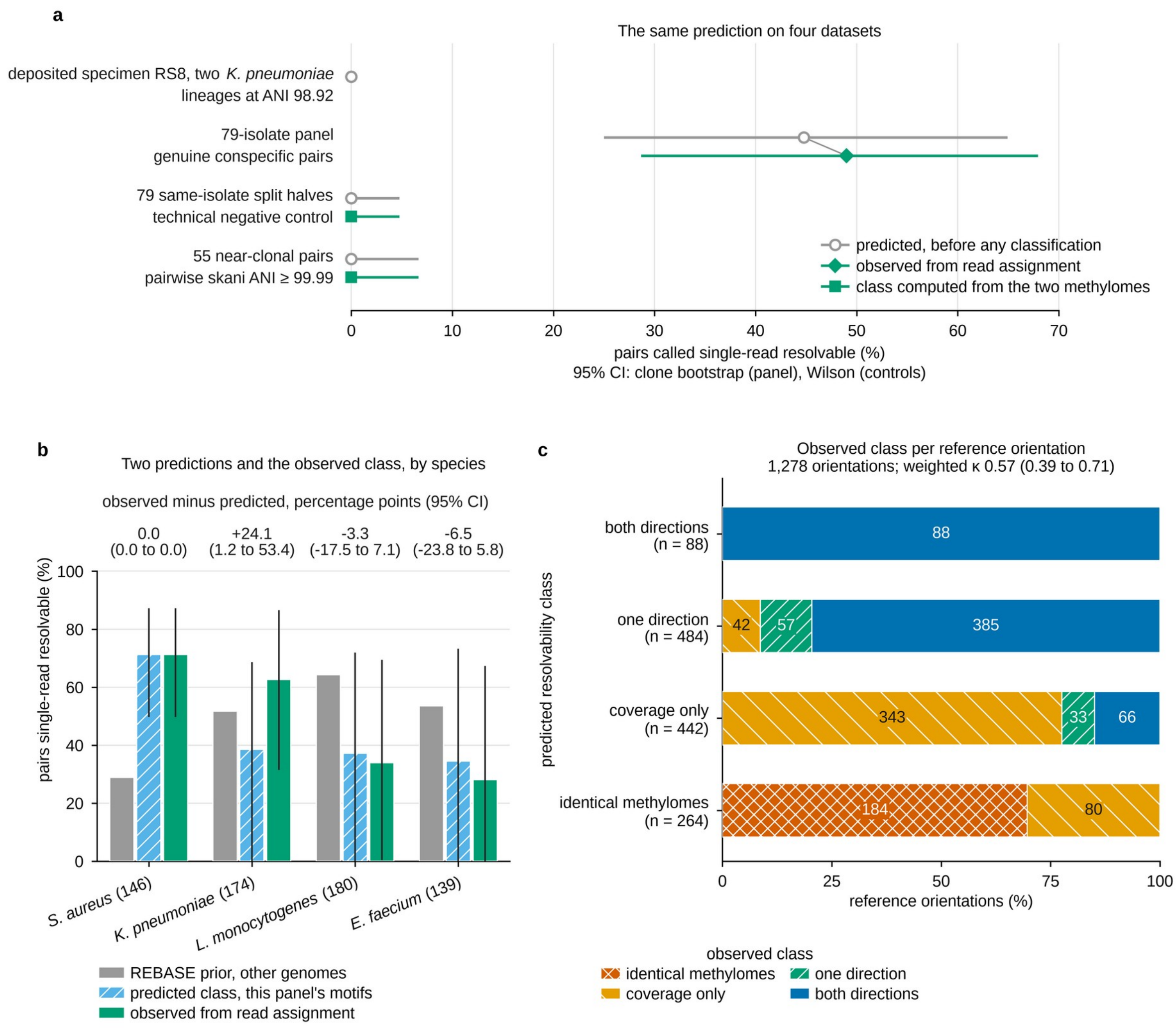


**Fig. S4.** The resolvability prediction tested on four datasets, by species and per reference orientation. (a) Pairs predicted single-read resolvable (circles) against, for the panel, the fraction observed from read assignment at default settings (diamond) and, for the two controls, the class computed from the two methylomes (squares), which is not a read-assignment outcome. Top to bottom: specimen RS8, two *Klebsiella pneumoniae* lineages at 98.92% ANI, predicted not resolvable (n = 1, no per-read outcome); the 639 genuine conspecific pairs of the 79-isolate panel, 44.8% predicted (242 one direction, 44 both directions) and 49.0% (313) observed, the class agreeing in 333 (52.1%; linearly weighted kappa 0.57, 95% CI 0.39 to 0.71); 79 same-isolate split halves, 0 of 79; 55 near-clonal pairs (ANI ≥ 99.99%), 0 of 55. The panel's intervals are clone bootstraps over 50 clones, because pairs share isolates; the controls' are Wilson intervals that treat units as independent. (b) By species, a REBASE prior from unrelated genomes, the class predicted from this panel's motifs and the observed class, with clone-bootstrap intervals (none on the prior). Observed minus predicted in percentage points (95% CI): *Staphylococcus aureus* 0.0 (0.0 to 0.0), *K. pneumoniae* +24.1 (1.2 to 53.4), *Listeria monocytogenes* −3.3 (−17.5 to 7.1) and *Enterococcus faecium* −6.5 (−23.8 to 5.8; 8 clones). (c) Observed class in each of the 1,278 reference orientations (each strain once the reference), by predicted class, under the same rule; the class agreed in 672 (52.6%; kappa 0.57, 0.39 to 0.71).

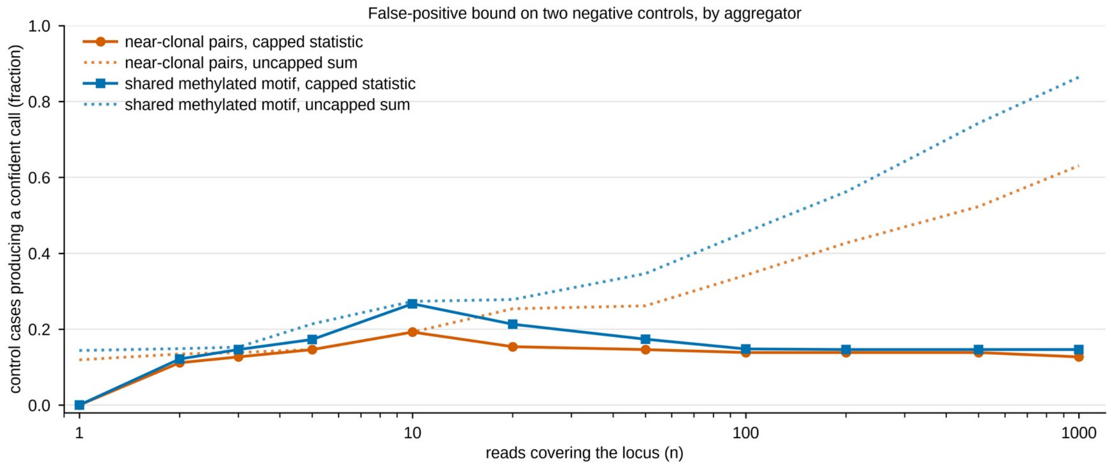


**Fig. S5.** Capping the locus score bounds false confident calls on two negative controls. Fraction of control cases producing a confident host call more than 1% of the time, against the number of reads covering the locus, under the locus score Λ = min(n, 10) × mean per-read evidence clipped at ±5, where n is the number of reads covering the locus (capped, solid), and under an uncapped sum of per-read evidence (dotted). The controls are near-clonal pairs, where only one strain is present so every confident call is false (n = 260 cases), and a shared methylated motif, a model that cannot discriminate (n = 1,244 cases). At a thousand reads the uncapped sum reached 0.631 and 0.864; the capped score converged to 0.127 and 0.146. Near-clonal false positives concentrate where no shared-motif control model can be built: 0.4286 of cases (n = 56) against 0.0000 where one exists (n = 204), at a locus score threshold of 10 (source data only). Measured at default settings, with reads the fidelity filter empties of calls counted at zero evidence.

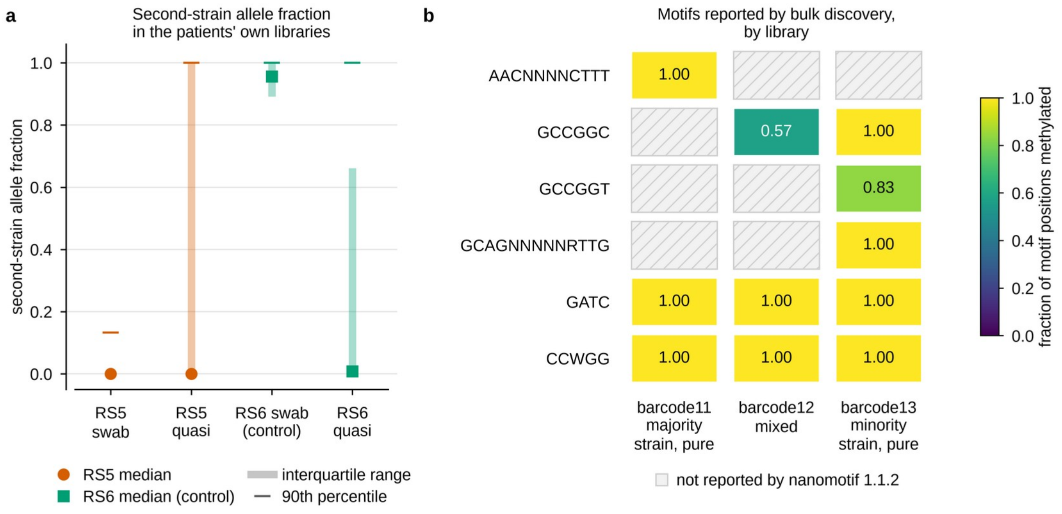


**Fig. S6.** The two-strain deposited library in detail. (a) Second-strain allele fraction at diagnostic sites in the two patients' own native metagenomes and quasi-metagenomes (the depositors' name, in ENA PRJEB108076, for a second library prepared from the same rectal swab): median (circle for RS5, square for the RS6 positive control), interquartile range (bar) and 90th percentile (tick), over 78,719 to 123,444 sites per library at median depths of 12× to 124×. The second strain of library RS5_barcode12 (patient RS5's deposited isolate library on barcode 12) is absent from patient RS5's own swab (median 0.000, 90th percentile 0.133), whereas the positive control from patient RS6 gives a median of 0.957. (b) Fraction of motif positions methylated for six motifs in the majority strain's pure library, the mixed library and the minority strain's pure library, as reported by bulk motif discovery (nanomotif 1.1.2); hatched tiles are motifs not reported for that library. Bulk discovery on the mixed library fails in both directions: AACNNNNNCTTT is reported for the pure majority library (1.000) but not for the mixture, and GCCGGC is reported for the mixture at 0.567 although it belongs to the minority strain (0.998 in its own library). The mechanism that placed two strains in one deposited library is undetermined.

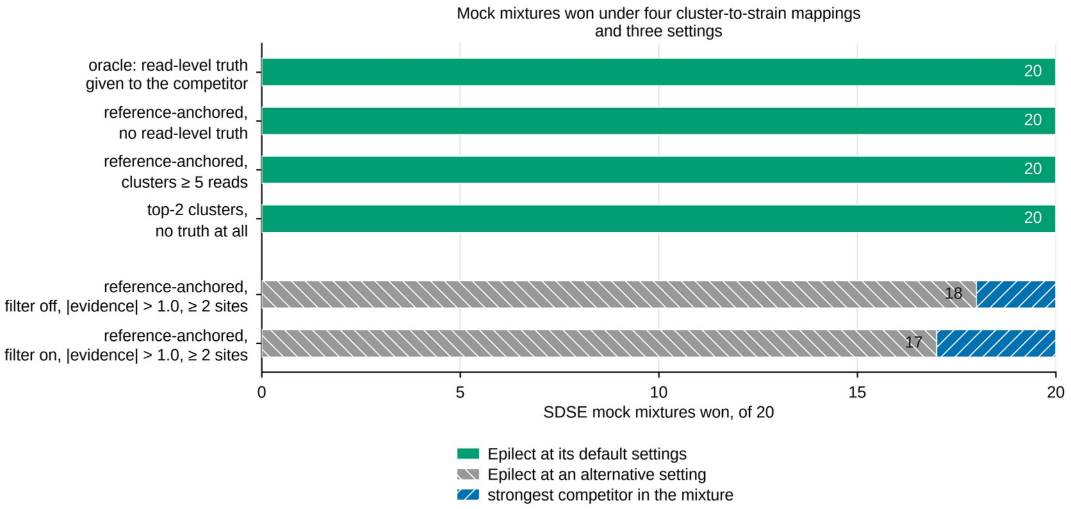


**Fig. S7.** At default settings Epilect wins all 20 SDSE mock mixtures under four rules for mapping a competitor's read clusters to strains. Number of the 20 mixtures of two *Streptococcus dysgalactiae* subsp. *equisimilis* (SDSE) isolates in which Epilect's per-read F1 (harmonic mean of precision and recall) over every offered read exceeds that of the strongest competitor in the same mixture. That every tool was scored over one offered read set is asserted rather than counted: all 20 runs record an identical scored-read source, but no mixture was recounted read by read. Mappings differ in the truth the competitor receives: top-2 clusters, none; reference-anchored mapping labels a cluster by the strain-private alleles its consensus carries, with and without a five-read minimum cluster size; the oracle mapping supplies read-level truth. At default settings (fidelity filter on, |evidence| > 1.1, at least one informative site) Epilect won 20 of 20 mixtures under every mapping. Hatched bars give two alternative settings on the reference-anchored mapping: 18 of 20 with the filter off at |evidence| > 1.0 and at least two informative sites, and 17 of 20 with the filter on at the same threshold. Competitors were Floria, Floria -p 2, devider, devider --sensitive, HairSplitter and Strainy; Strainy is not reproducible run to run because metaFlye pins no seed. These mixtures overstate the margin relative to the 704-mixture benchmark of Fig. 5.

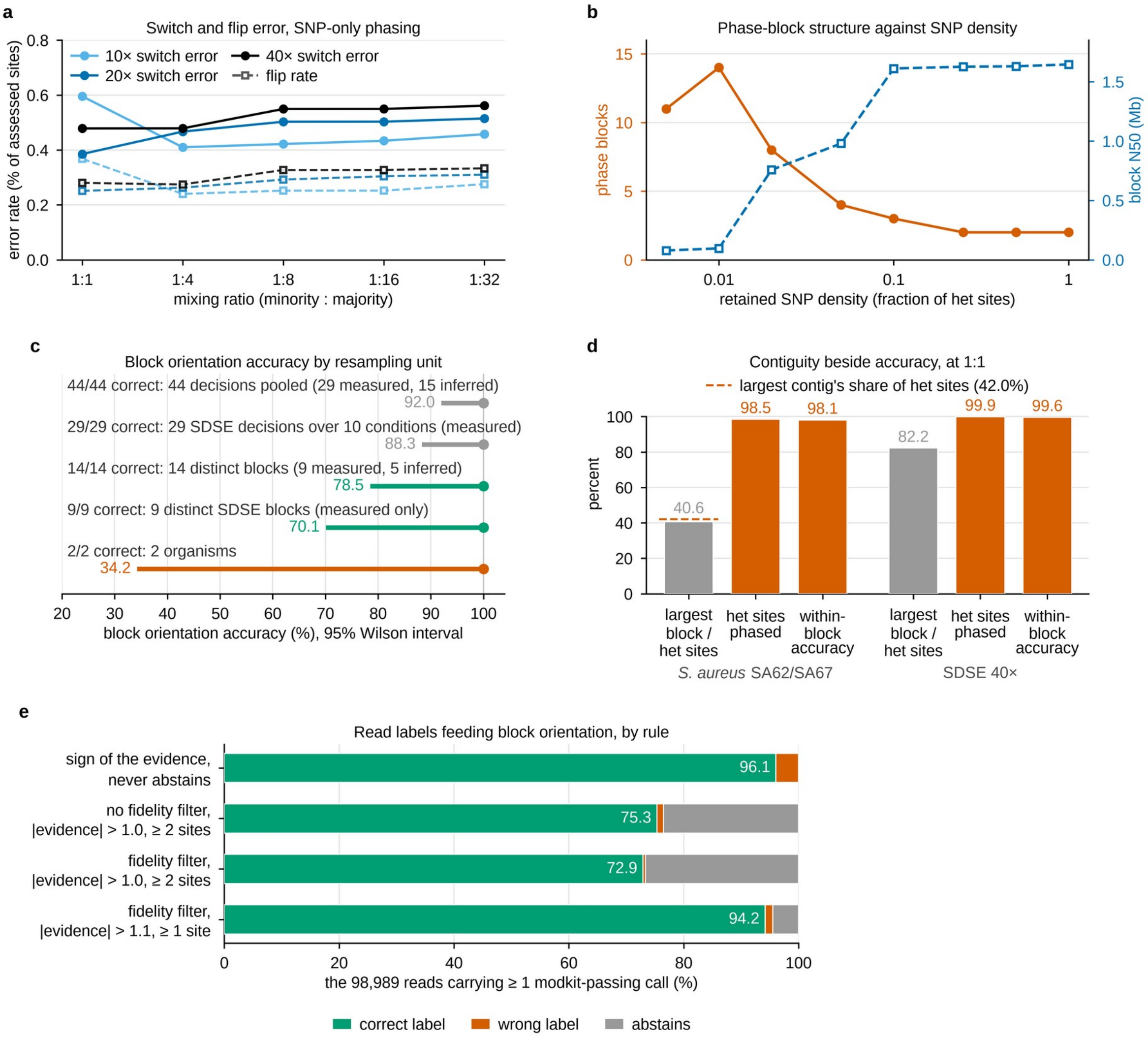


**Fig. S8.** Single-nucleotide polymorphism (SNP)-based phasing of the mock mixtures, and the methylation read labels that orient its blocks. (a) Switch error (solid) and flip rate (dashed) for read-backed SNP-only phasing of the mock mixtures of two *Streptococcus dysgalactiae* subsp. *equisimilis* (SDSE) isolates (Fig. S1) at three depths and five mixing ratios; over the 15 mixtures switch error stayed between 0.39% and 0.60% of assessed sites. (b) Phase blocks (left axis) and block N50 (right axis; the length at which blocks that long or longer cover half the phased sequence) as heterozygous (het) sites are removed. (c) Block orientation accuracy with 95% Wilson intervals under five choices of independent unit (grey: decisions; green: distinct blocks; vermilion: organisms); every decision was correct, and the lower bound is 92.0% over 44 pooled decisions, 78.5% over 14 distinct blocks and 34.2% over two organisms. (d) At 1:1, the largest block as a percentage of het sites, het sites phased and within-block accuracy, for *Staphylococcus aureus* SA62/SA67 (29,885 het sites) and SDSE at 40× (17,154); the dashed line is the largest contig's share of het sites (42.0%), which caps the first statistic. (e) Read labels under four rules, over the 98,989 pure control reads with at least one call passing modkit 0.6.4 filtering: the sign of the evidence labels 96.06% correctly; default settings label 94.18% correctly, with 1,330 errors and 4,434 abstentions.

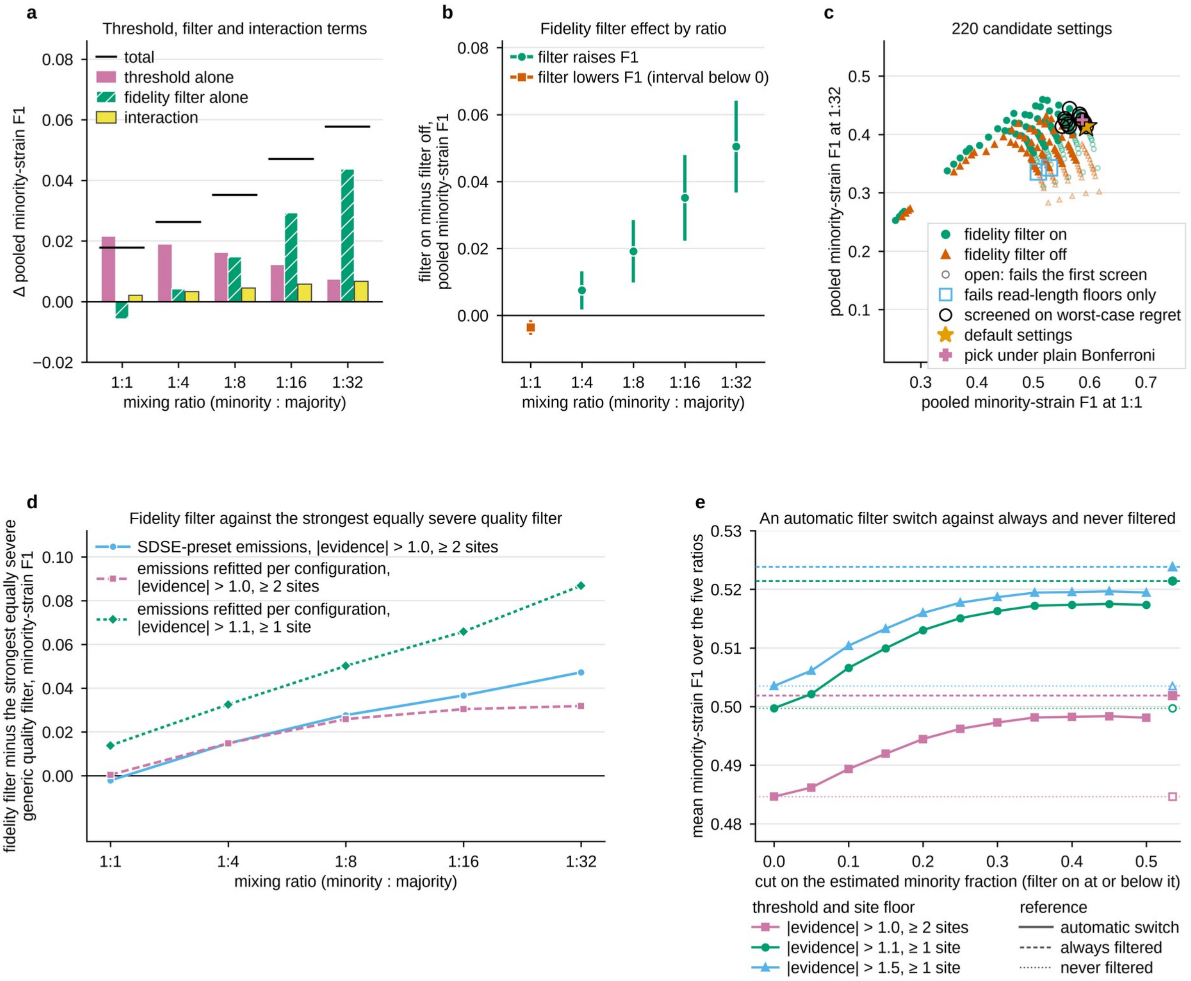


**Fig. S9.** Selection of the default settings. Panels a to c give pooled minority-strain F1 (harmonic mean of precision and recall), and panel e its mean over the five ratios, over 1,278 genuine non-near-clonal pairings (two strains, one the reference genome), each scored with either isolate as the minority (2,556 mixtures); panel d is the SDSE mock (Fig. S1), three mixtures per ratio. (a) Change in pooled F1 from the filter off at |evidence| > 1.0 and two sites, split into threshold, filter and interaction terms; the black rule is the total. (b) Effect of the fidelity filter alone at the default threshold, with paired leave-one-clone-out 95% intervals over 50 clones (circles where the filter raises F1, the square where it lowers it). (c) All 220 candidate settings, by pooled F1 at 1:1 against 1:32; open markers fail the first screen. Twelve matched or beat the default's worst-case regret and went to a second screen of 110 stratum-level tests per candidate (170 with six read-length bins, same pick; Table S56). Only the default passed under familywise control by the family's maximum-|z| null (critical value 3.205 for the 110 tests), Holm, Bonferroni at the effective number of tests or Benjamini-Hochberg; under plain Bonferroni three points pass and the pick would move to the rival marked +. (d) The fidelity filter minus the strongest equally severe generic quality filter, in three configurations; each point is a mean of three mixtures, without interval. (e) An automatic filter switch set by the estimated minority fraction; dashed lines are always filtered and dotted lines never filtered.
